# The benzoxazinoid breakdown product 5-Methoxy-2-Nitrophenol has a dual function in maize defense against herbivory

**DOI:** 10.1101/2023.10.10.561717

**Authors:** Bernardus C. J. Schimmel, Rocío Escobar-Bravo, Pierre Mateo, Cindy Chen, Gaétan Glauser, Vera Ogi, Lei Wang, Arletys Verdecia-Mogena, Christelle A. M. Robert, Matthias Erb

## Abstract

Catabolism of plant secondary metabolites can yield bioactive breakdown products. However, these compounds often remain poorly characterized. Here, we report on the discovery, biosynthesis, and biological relevance of 5-Methoxy-2-Nitrophenol (5M2NP), a secondary metabolite breakdown product which accumulates in damaged maize tissues. We used mutant plants, biochemical complementation, and metabolomic analyses to determine the biosynthetic origin of 5M2NP. Additionally, we assessed 5M2NP’s role in plant defense against herbivores. For this, we surveyed defense-associated responses (gene expression, phytohormones, volatile organic compounds) in wounded, 5M2NP-supplemented maize leaves, and performed bioassays with herbivorous insects. We found that 5M2NP accumulation upon tissue disruption is contingent upon a functional benzoxazinoid biosynthesis pathway. Labeling experiments demonstrated that 5M2NP is derived from DIMBOA. Physiological doses of exogenous 5M2NP increased the wound-induced expression of defense genes and emission of terpenoids. Additionally, 5M2NP exhibited antibiotic and antixenotic activities towards both generalist and specialist herbivores in nano-to micromolar quantities. We conclude that 5M2NP, which represents a novel class of plant-derived compounds, can act as a direct defense and a defense modulator. 5M2NP thus expands the functional repertoire of benzoxazinoids and likely contributes to their protective function against insect herbivores. The discovery of 5M2NP highlights the multifunctionality of plant secondary metabolites and their breakdown products.

## INTRODUCTION

Plants recognize herbivore attack by detecting various small molecules (Erb and Reymond, 2019; Snoeck et al., 2022). Collectively referred to as ‘danger signals’, these small molecules are commonly categorized in accordance with their origin: damage-associated molecular patterns (DAMPs) are derived from the plant itself, while herbivore-associated molecular patterns (HAMPs) are derived from herbivores (Snoeck et al., 2022; Tanaka and Heil, 2021). Plant perception of danger signals by cell-surface-localized receptors results in the activation of a signaling cascade, which involves ion and calcium fluxes, bursts of reactive oxygen species (ROS), activation of protein kinases, accumulation of phytohormones (primarily jasmonates and salicylic acid), and profound changes in gene expression (Couto and Zipfel, 2016; Schuman and Baldwin, 2016; Zhou and Zhang, 2020). Ultimately, it results in the production and/or activation of defensive molecules and metabolites, also known as secondary or specialized metabolites, which deter or otherwise harm the herbivore (Kant et al., 2015; Howe and Jander, 2008; Mithöfer and Boland, 2012; Unsicker et al., 2009; Erb and Reymond, 2019). Plant defenses can also act indirectly by increasing the recruitment and retention of natural enemies of the attacker, for example via emission of volatile organic compounds (VOCs), which serve as host-finding cues for parasitoids or predators of arthropod herbivores (Turlings and Erb, 2018; Unsicker et al., 2009). Besides in locally damaged tissues, plant defenses are also activated or primed in adjacent cells and more distal systemic tissues to prepare these for a potential imminent attack (Vlot et al., 2021; Mauch-Mani et al., 2017).

Plant secondary metabolites can be highly multifunctional (Erb and Kliebenstein, 2020; Weng et al., 2021; Neilson et al., 2013; Piasecka et al., 2015). Certain terpenes, flavonoids, green leaf volatiles (GLVs), glucosinolates and benzoxazinoids not only act as defenses, but also as regulators of growth and defense responses (Erb and Kliebenstein, 2020; Neilson et al., 2013). Multifunctionality has been well studied for glucosinolates and benzoxazinoids. Both classes of secondary metabolites are stored as stable glucosides and spatially separated from dedicated activating enzymes that cleave off the sugar molecule(s) (Frey et al., 2009; Halkier and Gershenzon, 2006). Wounding, for instance due to herbivory, mixes the glucosides with glucosidases yielding unstable and highly reactive aglucone intermediates that, depending on environmental conditions, break down into more stable compounds (Halkier and Gershenzon, 2006; Wouters et al., 2016b). Other plants have evolved similar wound-inducible two-component defense systems (Mithöfer and Boland, 2012; Kant et al., 2015; Morant et al., 2008). Hence, plant secondary metabolites may represent a major and reliable source of DAMPs and thereby facilitate the recognition of herbivory and pathogen infection (Erb and Kliebenstein, 2020). Most secondary metabolite catabolites and their functions, however, remain to be discovered.

Breakdown products of secondary metabolites can have distinct and specific functions or effects in plants. Both glucosinolates and benzoxazinoids, for instance, have been implicated in the herbivore or pathogen-induced deposition of callose at cell walls of attacked tissues, which serves as a physical barrier to small parasites. Arabidopsis (*Arabidopsis thaliana*) indole glucosinolate biosynthesis mutants are impaired in their callose deposition response upon treatment with Flg22 (Clay et al., 2009), a danger signal derived from bacterial flagellin. The response can be reinstated by supplementing the mutant plants with 4-methoxy-indol-3-ylmethylglucosinolate, but this requires the thioglucosidase PEN2, indicating that one or more hydrolysis products of 4-methoxy-indol-3-ylmethylglucosinolate function as DAMPs to regulate this defense response (Clay et al., 2009). Likewise, the maize (*Zea mays*) benzoxazinoid biosynthesis double mutant *bx1.igl* is unable to mount a full-scale callose response upon treatment with chitosan (Ahmad et al., 2011), a danger signal derived from fungal and arthropod chitin. Leaf infiltration with the benzoxazinoid DIMBOA, on the other hand, induces callose deposition in wild type maize (Ahmad et al., 2011) and wheat (*Triticum aestivum*) (Li et al., 2018). Glucosinolate breakdown products have further been shown to: i) inhibit root growth, presumably via competitive binding of auxin receptor TIR1 (Katz et al., 2015); ii) regulate stomatal aperture via ROS signaling (Zhao et al., 2008; Khokon et al., 2011; Hossain et al., 2013; Salehin et al., 2019), and; iii) elicit membrane depolarizations, calcium fluxes and jasmonate responses in distal leaves upon a local wounding event (Gao et al., 2023). Whether benzoxazinoid breakdown products also act as defense regulators is unknown, although it is likely given their abundance and reactivity.

Here, we identified 5-Methoxy-2-Nitrophenol (5M2NP) in damaged maize tissues, a compound that represents a hitherto unknown class of plant-derived metabolites. Based on its structural relatedness to DIMBOA and MBOA, we hypothesized that 5M2NP may be a benzoxazinoid-derivative. We analyzed several benzoxazinoid biosynthesis mutants for their 5M2NP content and performed biochemical complementation assays, including with labeled DIMBOA, to confirm this hypothesis. Since 5M2NP accumulates upon tissue disruption, we monitored gene expression changes, phytohormone concentrations and VOC emissions in maize leaves following wounding and 5M2NP-supplementation treatments to investigate if 5M2NP modulates defense responses. Finally, as benzoxazinoids can deter and hamper herbivores (Wouters et al., 2016a), we conducted a series of bioassays with leaf and root-feeding insects to explore 5M2NP’s impact on their performance and behavior. Together, this work demonstrates that 5M2NP has a dual function in maize defense against herbivory – it can act as a direct defense and as a defense modulator.

## MATERIALS AND METHODS

### Plants and insects

The following maize (*Zea mays* L.) genotypes were used: Delprim, B73, *bx1* (in the B73 background; Maag et al., 2016), *bx1.igl* (line 32R; Ahmad et al., 2011), W22, *bx1/W22* (Betsiashvili et al., 2015), F2, and the *brown-midrib* mutants *bm1*, *bm2*, *bm3* and *bm4* (in the F2 genetic background; Guillaumie et al., 2007; Barrière et al., 2004). Maize seeds were sown individually in plastic pots (height 11 cm, diameter 4 cm) in sand topped with commercial potting soil (Selmaterra; Bigler Samen AG, Thun, Switserland), unless indicated otherwise. Seedlings were germinated and grown in a glasshouse (day: 24 ± 2°C, 14-16 h light; night: 17 ± 1°C, 8-10 h dark; 40-60% relative humidity [RH]), including during experiments, unless indicated otherwise.

Eggs of *Spodoptera frugiperda* (J. E. Smith) and *Spodoptera littoralis* (Boisduval) were obtained from the University of Neuchâtel (Neuchâtel, Switzerland) and Syngenta Crop Protection AG (Stein, Switzerland), respectively. *Spodoptera* larvae were reared on a soy-wheat germ-based artificial diet (F9772; Frontier Agricultural Sciences, Newark, DE, USA) inside a climate room (24 ± 2°C; 14: 10 h, light: dark; 40-60% RH) until use. Eggs of *Diabrotica virgifera virgifera* (LeConte) and *Diabrotica balteata* (LeConte) were obtained from the USDA-ARS (Brookings, SD, USA) and Syngenta Crop Protection AG, respectively. Depending on the experiment, *Diabrotica* larvae were reared on W22 or *bx1/W22* seedlings grown in commercial potting soil, inside a climate chamber (24°C; 12: 12 h, light: dark; 40% RH), until use.

### 5M2NP identification and quantification

Two distinct methods were employed to determine (relative) amounts of 5M2NP in maize tissues. SPME-GC-MS, as outlined in Escobar-Bravo et al. (2022b), was used as the standard method. Samples were aliquoted from ground frozen homogenized plant material (roots: 90-110 mg, leaves: 10-20 mg) prior to analysis. Note that this method involves incubating individual samples at 50°C for 40 min to collect VOCs (Escobar-Bravo et al., 2022b). 5M2NP identification was based on similarity to NIST14 MS Library matches (NIST, Gaithersburg, MD, USA), plus comparisons with mass spectra and retention times of commercially available methoxy-nitrophenols (Supporting Information Fig. S1).

In a complementary approach, 5M2NP was extracted from mechanically damaged or undamaged control leaves of 14-d-old B73 seedlings (n = 4) with methanol and quantified by means of UHPLC-MS/MS, using an Acquity I-Class UPLC system (Waters Corp., Milford, MA, USA) coupled to a Xevo TQ-XS mass spectrometer (Waters) through an atmospheric pressure chemical ionization source (Methods S1). Prior to 5M2NP extraction, sample aliquots, containing *c*. 100 mg of ground frozen homogenized leaf material produced from each biological replicate, were subjected to a secondary treatment. That is, one aliquot was incubated at room temperature (RT) for 20 min, thus allowing the tissue to thaw. A second aliquot was incubated at 40°C for 20 min, while a third aliquot was kept at –80°C as a control.

For both methods, 5M2NP quantification in plant samples was based on an external concentration series of the synthetic standard.

### Root 5M2NP content of seedlings grown in soil versus soil-free conditions

W22 seeds were surface-sterilized with 5% bleach, rinsed three times with Milli-Q water, and then individually germinated and grown either in sand and soil or in a soil-free system, as in Hu et al. (2018). Crown roots were collected from 16-d-old seedlings, flash frozen in liquid nitrogen and stored at –80°C until analysis (n = 6).

### Supplementation of ground *bx1* tissue with DIMBOA or MBOA

Crown roots were collected from 23-d-old *bx1* seedlings, flash frozen in liquid nitrogen and stored at –80°C until use. Root powder from two plants was pooled to form one biological replicate. From each biological replicate, a 100 mg aliquot was supplemented with 100 µl of 0.5 mg DIMBOA ml^-1^ Milli-Q water, vortexed for 2 min, and incubated at RT until analysis. As a control, another 100 mg aliquot of frozen root powder from the same biological replicate was supplemented with 100 µl Milli-Q water and treated and analyzed in the same way. This experiment was repeated with MBOA instead of DIMBOA, using the same metabolite concentration and plant material from the same set of samples. 5M2NP was quantified after *c.* 0.1 h, 1.5 h, 3.0 h, 4.6 h and 6.1 h of incubation. Amounts of DIMBOA and MBOA used for *bx1* complementation are consistent with the DIMBOA concentration found in crown roots of B73 maize (Machado et al., 2021). Each complementation experiment was performed in five blocks (experimental replicates) in time, in total with five biological replicates per treatment and incubation period (n = 5). As additional controls, a vial with 100 µl of 0.5 mg DIMBOA ml^-1^ Milli-Q water and a vial with 100 µl Milli-Q water were included in each experimental replicate. Chemical solutions were freshly prepared for each experiment.

Complementation of *bx1* plant material (crown roots and leaves) with DIMBOA-*d*3 was performed as above, with minor modifications. Root and leaf powder was obtained from individual 16-d-old *bx1* seedlings. Each sample aliquot of 100 mg frozen plant tissue was supplemented with 100 µl of 0.5 mg DIMBOA-*d*_3_ ml^-1^ Milli-Q water, vortexed for 2 min, and incubated for 1.5 h at RT prior to analysis. Aliquots of the DIMBOA-*d*_3_ solution were stored at –20°C until use. This experiment was limited to two biological replicates per treatment and tissue type (n = 2), because we detected quantitatively similar synthesis of 5M2NP-*d*_3_ from DIMBOA-*d*_3_ in the presence and absence of plant material (see Results section).

### Pharmacological treatments

W22 seeds were surface-sterilized, germinated and grown in a soil-free system as described earlier. Seedlings were supplemented every 3 d alternately with, depending on humidity of the system, 3-5 ml of a chemical solution (on d 9, 15, 21) or a nutrient solution (on d 12 and 18). Chemical solutions contained 1 mM of either shikimic acid, phenylalanine, caffeic acid, or piperonylic acid. Milli-Q water (1% v/v ethanol) was used as a control. Crown roots were collected from 23-d-old seedlings, flash frozen in liquid nitrogen and stored at –80°C until analysis (n = 10).

### Benzoxazinoid profiling

Benzoxazinoids were extracted from crown roots of 16-d-old *bm* mutants and F2 seedlings (n = 5) and quantified by means of UHPLC-MS, using an Acquity I-Class UPLC system (Waters) coupled to a UV detector (PDA eλ; Waters) and QDa mass spectrometer equipped with an electrospray source (Waters), as in Machado et al. (2021).

### Wound and defense-associated responses in 5M2NP-supplemented damaged maize leaves

The third leaf of 16-d-old *bx1* seedlings was wounded four times, perpendicular to but not crossing the midrib, using a hemostat. Wounds were inflicted *c.* 5 cm from the leaf-base and immediately supplemented with 20 µl of a 5M2NP solution (7.5 ng 5M2NP µl^-1^) or Milli-Q water (0.75% v/v dichloromethane) as a control. Each wound received *c.* 5 µl treatment solution, divided over the abaxial and adaxial sides. The amount of 5M2NP for *bx1* complementation was determined based on the average 5M2NP concentration in leaves of wild type (WT) maize (1.2 ng mg^-1^ fresh weight [FW]; Fig. S2a) and the average mass of the targeted leaf section (*c.* 125 mg). For local gene expression analysis, 5 cm long leaf sections centered around the wounds were cut out at 0.5 h, 1 h, 4 h and 8 h after treatments, flash frozen in liquid nitrogen and stored at –80°C until we isolated the RNA (n = 8). Corresponding leaf sections from eight untreated plants sampled at the beginning of the experiment (0 h) were used as controls.

For local phytohormone analysis, plants were treated and sampled as described in the previous section, except for the 8 h time point, which was excluded. Wound + water-treated B73 plants were included as additional controls. Samples were stored at –80°C until we extracted the hormones (n = 12).

For the analysis of VOCs emitted from wound + 5M2NP or wound + water-treated plants, 16-d-old *bx1* seedlings, placed individually in closed cylindrical glass chambers (height 45 cm, diameter 12 cm), were treated as described above and, starting 10 min after treatments, headspace volatiles were analyzed in real-time by PTR-TOF-MS every 30 min (for 56 s) over a period of 12 h (n = 11). Plants were transferred to glass chambers 1 d before the start of the experiment. Baseline (control) emission levels of plant VOCs were determined by analyzing headspace volatiles over a period of 1.5 h prior to treatments. FW of the aboveground plant material was recorded 24 h after treatments.

### Gene expression analysis

Transcript abundances of selected maize genes in sampled leaf sections were determined by qRT-PCR, using a 384-well Applied Biosystems QuantStudio 5 Real-Time PCR system (Thermo Fisher Scientific) following the procedures described by Schimmel et al. (2017) and Escobar-Bravo et al. (2022b), with minor modifications (Methods S2). *Ubiquitin* was used as reference gene. Target genes, their identifiers, primer sequences and references are listed in Table S1.

### Phytohormone analysis

Phytohormone concentrations in sampled leaf sections were determined by UHPLC-MS/MS, using an Acquity UPLC system (Waters) connected to a QTRAP 6500+ mass spectrometer (SCIEX, Framingham, MA, USA) following Glauser et al. (2014) and Schimmel et al. (2017), with minor modifications (Methods S3).

### Analysis of plant volatiles

Plant headspace VOCs were sampled non-invasively and analyzed using a high-throughput phenotyping platform with integrated Vocus PTR-TOF-MS autosampler (Tofwerk AG, Thun, Switzerland) and Tofware software v3.2.2 (Tofwerk AG), as described in Escobar-Bravo et al. (2022a), in a dedicated room under controlled environmental conditions (day: 28°C, 14 h light, 375 ± 25 µmol m^-2^ s^-1^; night: 20°C, 10 h dark; 30-40% RH).

### Herbivore preference assays

Petri dish-based choice assays with *S. littoralis* and *S. frugiperda* larvae were adapted from Glauser et al. (2011) and Rharrabe et al. (2014). In short, we monitored the distribution over time of groups of five caterpillars given a choice between two differentially treated artificial diet (AD) cubes: one supplemented with a 5M2NP solution (1.0 or 2.5 ng 5M2NP mg^-1^ AD) the other with Milli-Q water (0.9% v/v ethanol) as a control (Methods S4; n = 20). The amounts of 5M2NP for AD complementation were determined based on average and high 5M2NP concentrations in maize leaves (Fig. S2a).

Choice assays with *D. balteata* and *D. v. virgifera* larvae were adapted from Huang et al. (2017). We recorded the number of larvae rejecting a host plant, following the release of a group of five rootworms at the entrance of a 6.5 cm long glass tube that provided access to the rhizosphere of a *bx1* seedling, which had been supplemented with 5 ml of either a 5M2NP solution (5 or 25 ng 5M2NP ml^-1^) or autoclaved tap water (0.0025% v/v ethanol) as a control (Methods S5; n = 37 – 58). The amounts of 5M2NP for rhizosphere complementation were determined based on the average 5M2NP concentration in crown roots of WT maize (*c*. 0.1 ng mg^-1^ FW; Fig. S2a), a conservative estimate of *D. v. virgifera*’s root consumption (250 mg d^-1^ by five larvae; Zhang et al., 2019), and the average mass of the total root system of 16-d-old *bx1* seedlings (*c.* 1100 mg).

### Herbivore performance assays

No-choice assays with *S. littoralis* and *S. frugiperda* larvae were modified from Glauser et al. (2011). Briefly, we monitored the weight over time of individual caterpillars feeding on AD supplemented with either a 5M2NP solution at physiological doses (1.0 or 2.5 ng 5M2NP mg^-1^ AD) or Milli-Q water (0.9% v/v ethanol) as a control (Methods S6; n = 30).

Performance assays with *D. balteata* and *D. v. virgifera* were based on methods in Erb et al. (2011). Here, we determined the average weight gain of five *Diabrotica* larvae feeding for 4 d from *bx1* plants supplemented with either a 5M2NP solution at physiological doses (2.5, 25 or 125 ng 5M2NP d^-1^) or autoclaved tap water (0.0025% v/v ethanol) as a control (Methods S7; n = 12).

### Data analysis

All statistical analyses were performed using R software, v4.1.2 (R Core Team, 2021). Effects of plant genotype, plant tissue, treatment, time since treatment, and/or their interaction(s) on concentrations of metabolites (5M2NP, benzoxazinoids, phytohormones, VOCs), gene transcript abundances, and herbivore weight gain were assessed using linear models or linear mixed-effects models (LMMs). Effects of plant or AD treatment on herbivore preference (over time) were assessed using generalized linear mixed-effects models (GLMMs) with a Poisson distribution (Hu et al., 2018) and/or one-sample *t*-tests. When applicable, experimental replicate was included in the model as random factor. Fitted models were subjected to analysis of variance (ANOVA, Type II), repeated measures analysis of variance (RM-ANOVA, Type I) or analysis of deviance (Type II Wald chi-square), followed by estimated marginal means (EMMeans)-based pairwise *post hoc* tests with Tukey adjustment for multiple comparisons. Models were checked by inspecting the behavior and distribution of model residuals (Crawley, 2013). When necessary, data were transformed (i.e., log2, log10, ln, or sqrt) to correct for heteroscedasticity or non-normality of errors. When this was ineffective, nonparametric models/methods were used, including the aligned rank transform (ART) procedure (Wobbrock et al., 2011). Specifics are provided in the figure legends. Linear correlations (Pearson) were calculated between the concentrations of 5M2NP and individual benzoxazinoids. Data handling, analyses and visualizations were conducted using the following R packages: ‘ARTool’ (Kay et al., 2021), ‘car’ (Fox and Weisberg, 2019), ‘dplyr’ (Wickham et al., 2021), ‘emmeans’ (Lenth, 2021), ‘ggplot2’ (Wickham, 2016), ‘ggpubr’ (Kassambara, 2020), ‘Hmisc’ (Harrel Jr, 2021), ‘lme4’ (Bates et al., 2015), ‘lmerTest’ (Kuznetsova et al., 2017), ‘officer’ (Gohel, 2021) and ‘rvg’ (Gohel, 2020).

## RESULTS

### Identification of 5-Methoxy-2-Nitrophenol (5M2NP) from maize

We previously detected an unknown nitrophenol (*m/z* 139) as a compound of interest in an untargeted SPME-GC-MS-based metabolomic analysis of ground roots from maize plants (Huang et al., 2017). Subsequent analyses suggested that this unknown nitrophenol represented a fragment of a methoxy-nitrophenol (C_7_H_7_NO_4_; *m/z* 169). Using synthetic standards, the methoxy-nitrophenol was identified as 5M2NP (Fig. 1a and S1). 5M2NP was already known as synthetic chemical (John et al., 1975; Tokiwa and Ohnishi, 1986), but not yet as plant-derived substance. 5M2NP accumulated in both disrupted root and leaf tissues, but with a tenfold higher concentration in leaves than in crown roots and the lowest concentration in primary roots (Fig. S2a). 5M2NP concentrations were not significantly altered upon leaf infestation with *S. littoralis* or *S. frugiperda* larvae, or upon root infestation with *D. balteata* or *D. v. virgifera* larvae (Fig. S2).

**Figure 1:**
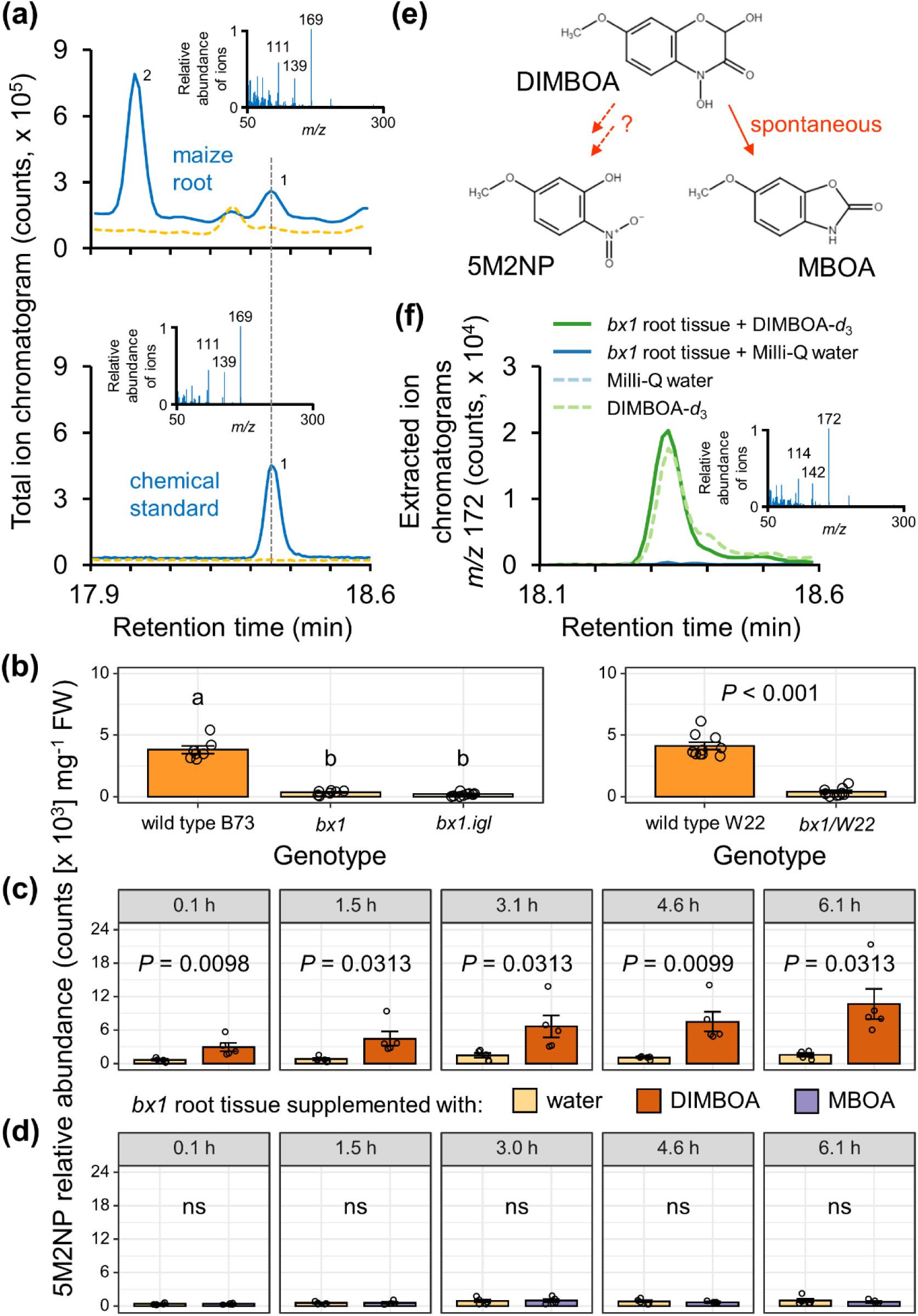
Identification and biosynthesis of 5-Methoxy-2-Nitrophenol (5M2NP). (a) Representative GC-MS total ion chromatograms of volatiles emanating from *c*. 100 mg ground crown roots of 21-d-old wild type maize plants (*Zea may*s cv. Delprim; top) or 2 ng of 5M2NP chemical standard (bottom), both in blue, and their respective controls in orange. Numbered peaks correspond to 5M2NP (#1) and (*E*)-ß-caryophyllene (#2). Insets depict reference electron ionization mass spectra of the respective 5M2NP peaks and include the mass-to-charge ratio (*m/z*) of the three most abundant ions. (b) Compared to wild type maize, the 5M2NP concentration is greatly reduced in *bx1* mutants, which do not have a functional BX1 enzyme and therefore accumulate only minor amounts of benzoxazinoids. Bars indicate mean (± SEM) amounts of 5M2NP in crown roots of 16-d-old *bx1* single (*bx1* or *bx1/W22*) and double (*bx1.igl*) mutants along with their corresponding wild types B73 or W22 (n = 7 – 10). Different letters above the bars in the panel on the left indicate statistically significant differences between genotypes as determined by a one-way analysis of variance (ANOVA), followed by estimated marginal means *post hoc* tests (Tukey-adjusted *P* ≤ 0.05). The *P*-value in the panel on the right represents the outcome of a Wilcoxon rank-sum test. (c, d) Supplementation of ground root tissue with DIMBOA, but not MBOA, restores the low-5M2NP phenotype of *bx1* mutants. Bars indicate mean (± SEM) amounts of 5M2NP emanating from *c*. 100 mg ground crown roots of 23-d-old *bx1* mutants supplemented with either 0.5 mg DIMBOA g^-1^ FW, 0.5 mg MBOA g^-1^ FW, or Milli-Q water as a control, after different incubation periods at room temperature (n = 5). Independent sets of samples were used for each incubation period. The *P*-values represent the outcome of either a one-tailed paired sample *t*-test or a one-tailed Wilcoxon signed-rank test (ns, not significant; *P* > 0.05). (e) Simplified schematic overview of DIMBOA catabolism, yielding either 5M2NP or MBOA. (f) Representative GC-MS extracted ion chromatograms of a diagnostic electron ionization ion (*m/z* 172) of deuterium (*d*_3_)-labeled 5M2NP, tentatively identified in vials loaded with 50 µg DIMBOA-*d*_3_, either or not in presence of *c*. 100 mg ground crown roots of 16-d-old *bx1* mutants and incubated for 1.5 h at room temperature, along with their respective controls. Inset: reference electron ionization mass spectrum of the putative 5M2NP-*d*_3_ peak from the DIMBOA-*d*_3_ sample. The mass-to-charge ratio of the three most abundant ions is included. FW, fresh weight

Next, we used UHPLC-MS/MS as an orthogonal analytical approach to confirm 5M2NP as a plant-derived metabolite (as opposed to a breakdown product that forms specifically in the context of SPME-GC-MS-based analyses). While no 5M2NP was detected in fresh extracts of frozen leaf material, a clear 5M2NP peak appeared in the chromatogram once the ground leaf material was incubated at RT for 20 min prior to sample extraction and analysis by UHPLC-MS/MS (Fig. S3a). Post-harvest incubation at a higher temperature, i.e. 40°C for 20 min, did not further increase the recorded 5M2NP concentration, nor did mechanical wounding of the leaf before harvest (Fig. S3b).

Our results demonstrate that 5M2NP accumulates in damaged maize tissues. As methoxy-nitrophenols have, to the best of our knowledge, not been described as plant-derived compounds, we set out to characterize the biosynthesis and biological function(s) of 5M2NP. Since the SPME-GC-MS-based method was more sensitive, it was used for all subsequent experiments.

### 5M2NP is derived from the benzoxazinoid DIMBOA

We performed a series of experiments to test hypotheses regarding the biosynthesis of 5M2NP. First, to assess the possible involvement of soil microbes in 5M2NP production, we quantified 5M2NP in roots of maize seedlings grown from surface-sterilized seeds in soil-free conditions. This environment excludes soil microbes but is not entirely sterile. The crown root 5M2NP content of these plants was not different from that of plants grown in sand and soil (Fig. S4), suggesting that 5M2NP is produced by maize itself rather than by soil microbes.

Since 5M2NP shares structural similarities with benzoxazinoids, such as DIMBOA, we hypothesized that 5M2NP is a benzoxazinoid derivative. Consistent with this hypothesis, we found that 5M2NP levels were reduced by 91 – 98% in roots and leaves of maize *bx1* mutants (Fig. 1b and S5), which do not have a functional BX1 enzyme and therefore accumulate only residual amounts of benzoxazinoids (Frey et al., 1997; Maag et al., 2016; Machado et al., 2021; Köhler et al., 2015). The low-5M2NP phenotype of *bx1* mutants was rescued by supplementing ground *bx1* root tissue with DIMBOA at a physiologically relevant dose (Fig. 1c; overall treatment effect: Wilcoxon signed-rank test, *P* = 5.96 x 10^-8^). By contrast, we did not detect increased amounts of 5M2NP in *bx1* root tissue supplemented with MBOA (Fig. 1d; overall treatment effect: Wilcoxon signed-rank test, *P* = 0.71). These results indicate that 5M2NP is derived from DIMBOA (Fig. 1e). An experiment with deuterium-labeled DIMBOA (DIMBOA-*d*3) confirmed this hypothesis, as DIMBOA-*d*_3_ was directly converted into 5M2NP-*d*_3_ (Fig. 1f). While only minor amounts of non-labeled DIMBOA were converted into 5M2NP in the absence of plant material, i.e. at levels similar to or less than those found in *bx1* root tissue supplemented with Milli-Q water (Fig. S6), surprisingly, the conversion rate of DIMBOA-*d*_3_ into 5M2NP-*d*_3_ was quantitatively similar in the presence or absence of plant material (Fig. 1f), suggesting that enzymatic action is not strictly required for the latter reaction.

Besides the benzoxazinoid pathway, numerous low molecular weight aromatic compounds are produced via the conserved phenylpropanoid pathway (Vogt, 2010). Products and intermediates of the phenylpropanoid pathway can also modulate the biosynthesis and transport of metabolites from other pathways (Yin et al., 2014; Kurepa et al., 2018; Kim et al., 2020). To examine the extent to which the phenylpropanoid pathway contributes to 5M2NP biosynthesis, we subjected WT plants to pharmacological treatments and performed chemical analyses of *brown-midrib* (*bm*) mutants. Repeated supplementation of maize seedlings with precursors (shikimic acid, phenylalanine) or an intermediate (caffeic acid) of the phenylpropanoid pathway did not significantly alter crown root 5M2NP levels (Fig. S7). Neither did supplementation with piperonylic acid, which is an inhibitor of cinnamate 4-hydroxylase (Schalk et al., 1998), the second key enzyme of the phenylpropanoid pathway. Likewise, the concentration of 5M2NP in crown roots of *bm1*, *bm3* and *bm4* seedlings, which have known mutations in other enzymes of the phenylpropanoid pathway (Halpin et al., 1998; Vignols et al., 1995; Li et al., 2015a), was similar to that of their WT parent, F2. In *bm2* roots, however, the 5M2NP concentration was about 30% lower (Fig. S8). The *bm2*-encoded methylenetetrahydrofolate reductase contributes to production of the methyl donor S-adenosyl-L-methionine (Tang et al., 2014). We subsequently used the same samples to profile the root benzoxazinoid content of the *bm* mutants (Fig. S8). A Pearson correlation analysis revealed that in *bm2*, but not in the other mutants nor in F2, concentrations of 5M2NP and one or more benzoxazinoids were significantly correlated. Specifically, we found linear correlations between 5M2NP and the R^3^-methoxylated (Wouters et al., 2016b) compounds HM_2_BOA-Glc (*R*^2^ = 0.92, *P* = 0.03), DIM_2_BOA-Glc (*R*^2^ = 0.90, *P* = 0.04) and HDM_2_BOA-Glc (*R*^2^ = 0.90, *P* = 0.04). Together, this suggests the phenylpropanoid pathway does not play a major role in 5M2NP biosynthesis but may act indirectly by modulating benzoxazinoid biosynthesis via metabolic crosstalk.

### 5M2NP promotes defense activation in wounded leaves

Considering that 5M2NP is derived from DIMBOA (Fig. 1), which is produced from DIMBOA-Glc upon tissue disruption, and that benzoxazinoids protect plants against arthropod herbivores in diverse ways (Wouters et al., 2016b; Erb and Kliebenstein, 2020), we explored 5M2NP’s role in plant defense. First, to investigate whether 5M2NP functions as a DAMP, we monitored molecular markers of wound and defense responses (gene transcripts, phytohormones, volatiles) over time in/from leaves with wounds supplemented with a physiological dose of 5M2NP. We used *bx1* mutants to minimize possible confounding effects of endogenous 5M2NP and benzoxazinoids.

As expected, tissue damage itself strongly induced expression of selected maize defense genes (Fig. 2; factor Time). Moreover, wound-induced transcript levels of half of these genes were higher following supplementation of wounds with 5M2NP (Fig. 2; factor Treatment). Compared to the wound + water control treatment, expression of genes involved in the biosynthesis of jasmonates (*LOX10*, *AOS1*; Christensen et al., 2013; Borrego and Kolomiets, 2016), proteinase inhibitors (*SerPIN*, *Cyst/CC9*; Ton et al., 2007; van der Linde et al., 2012), benzoxazinoids (*Bx1*; Frey et al., 1997), and the homoterpenes DMNT and TMTT (*CYP92C5*; Richter et al., 2016) was enhanced at one or more time points following the wound + 5M2NP treatment. A similar trend was observed for jasmonate biosynthesis gene *AOC1* (Borrego and Kolomiets, 2016). By contrast, the wound-induced expression of *Bx10/11* was dampened by 5M2NP supplementation. *Bx10* and *Bx11* code for *O*-methyltransferases which convert DIMBOA-Glc into HDMBOA-Glc (Meihls et al., 2013). Despite initial enhancement, dampening was later on also detected for *LOX10* and *CYP92C5*. We found no effect of 5M2NP supplementation on transcript levels of *TPS10*, *IGL* and *Bx14*, whose encoded enzymes synthesize sesquiterpenes (Schnee et al., 2006), volatile indole (Frey et al., 2000) and HDM_2_BOA-Glc (Handrick et al., 2016), respectively.

**Figure 2:**
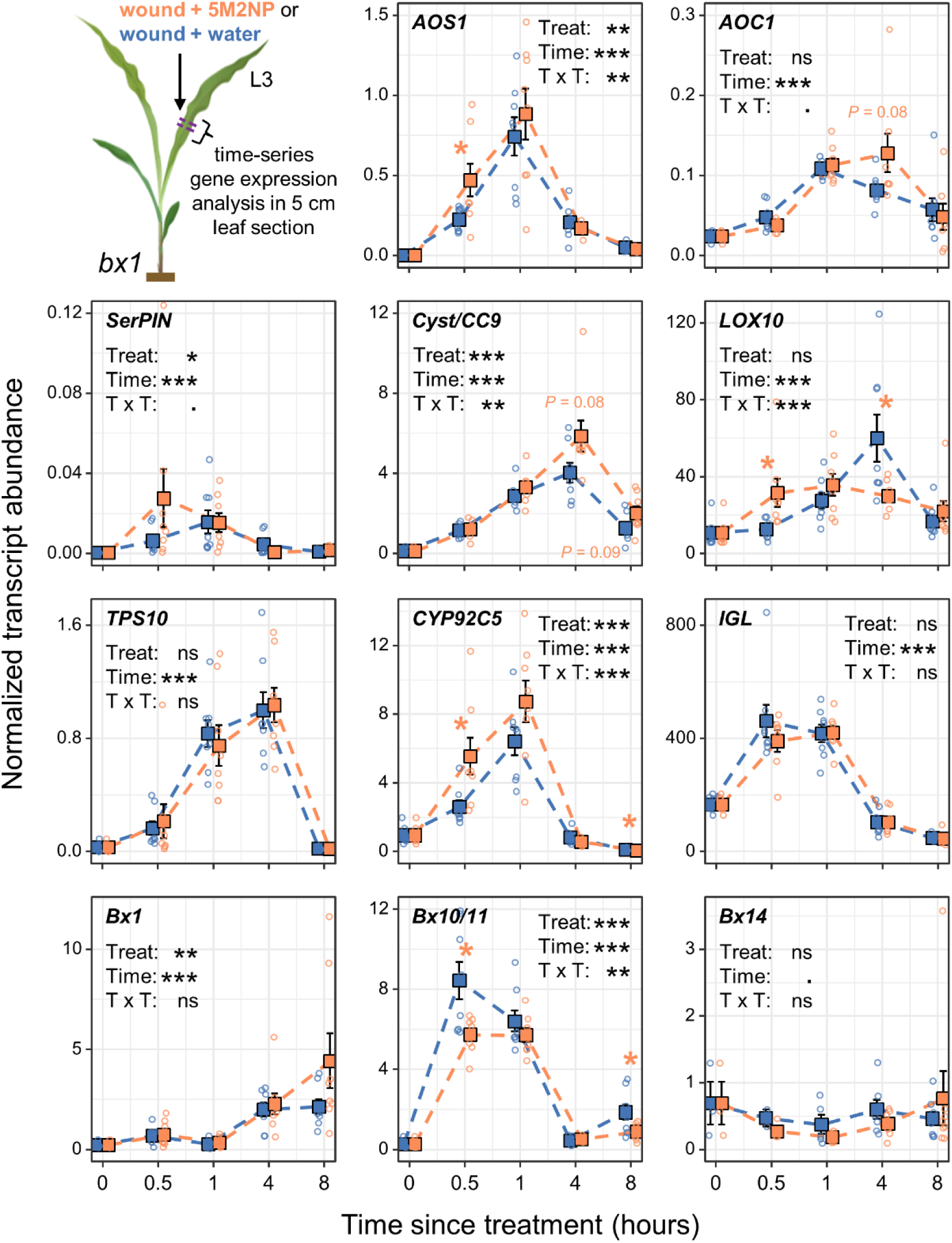
At a physiologically relevant dose, 5-Methoxy-2-Nitrophenol (5M2NP) modulates the expression of defense genes in wounded leaf tissue. The figure shows local, time-resolved expression profiles of wound and defense-associated genes in leaves of maize (*Zea mays*) *bx1* mutants following either a wound + 5M2NP treatment (in orange) or a wound + water treatment (in blue). The third leaf (L3) of 16-d-old *bx1* seedlings was wounded four times, perpendicular to but not crossing the midrib, using a hemostat. Wounds were immediately supplemented with 20 µl of a 5M2NP solution (7.5 ng 5M2NP µl^-1^) or Milli-Q water (0.75% v/v dichloromethane) as a control. At 0.5 h, 1 h, 4 h and 8 h after these treatments, 5 cm long leaf sections centered around the wounds were cut out for RNA isolation. Corresponding leaf sections from untreated plants sampled at the beginning of the experiment (0 h) were used as controls for both treatments. Squares depict mean (± SEM) normalized transcript abundances of the wounding/jasmonic acid marker genes *AOS1*, *AOC1*, *SerPIN*, *Cyst/CC9*, *LOX10*; volatile biosynthesis genes *TPS10*, *CYP92C5*, *IGL*; and benzoxazinoid biosynthesis genes *Bx1*, *Bx10/11*, *Bx14*, for each treatment, at indicated time points (n = 8; but note that gene transcripts were not consistently detected in all biological replicates, e.g. at 0 h). Transcript abundances were normalized to *Ubiquitin*. Black asterisks denote statistically significant effects of plant treatment (Treat), time since treatment (Time) and/or their interaction (T x T) on the expression level of each gene, as determined by a multifactorial analysis of variance on aligned rank-transformed data (ANOVA-ART): *** for *P* < 0.001; ** for *P* < 0.01; * for *P* < 0.05;. for *P* < 0.1.; ns, not significant for *P* > 0.1. Orange asterisks indicate statistically significant differences between treatments at individual time points according to an independent two-sample Student’s *t*-test, Welch’s *t*-test or Wilcoxon rank-sum test (*P* ≤ 0.05). Gene names are listed in Table S1.

Given the 5M2NP-augmented induction of jasmonate biosynthesis genes in wounded leaves (Fig. 2), we determined local concentrations of jasmonates and other stress-associated hormones in a separate experiment. Wound + water-treated B73 plants were included as additional controls. Within 0.5 h after wounding, amounts of 12-oxo-phytodienoic acid (OPDA), jasmonic acid (JA), as well as the main bioactive conjugate, JA-Ile, strongly increased, before gradually decreasing towards baseline levels (Fig. 3). Concentrations of salicylic acid (SA), abscisic acid (ABA) and indole-3-acetic acid (IAA; the predominant auxin) were also elevated in response to wounding, albeit more modestly and in the case of IAA only in B73. Two minor 5M2NP-mediated effects were discernible. First, IAA levels were significantly dependent on the interaction between the factors Treatment and Time, owing to intermediate – yet not consistently statistically different – concentrations found in 5M2NP-treated *bx1* wound sites at 1 h and 4 h. Second, 4 h after treatments, OPDA levels were lower in *bx1* wound sites treated with 5M2NP than with water. Of note, clear differences between genotypes were detected for most hormones at one or more time points. Concentrations of JA and JA-Ile were *c*. 2-fold higher in unwounded *bx1* leaves than in their WT counterparts, whereas SA followed the opposite pattern (Fig. 3). Overall, these results reveal no major contribution of 5M2NP to the wound-induced local accumulation of stress hormones.

**Figure 3:**
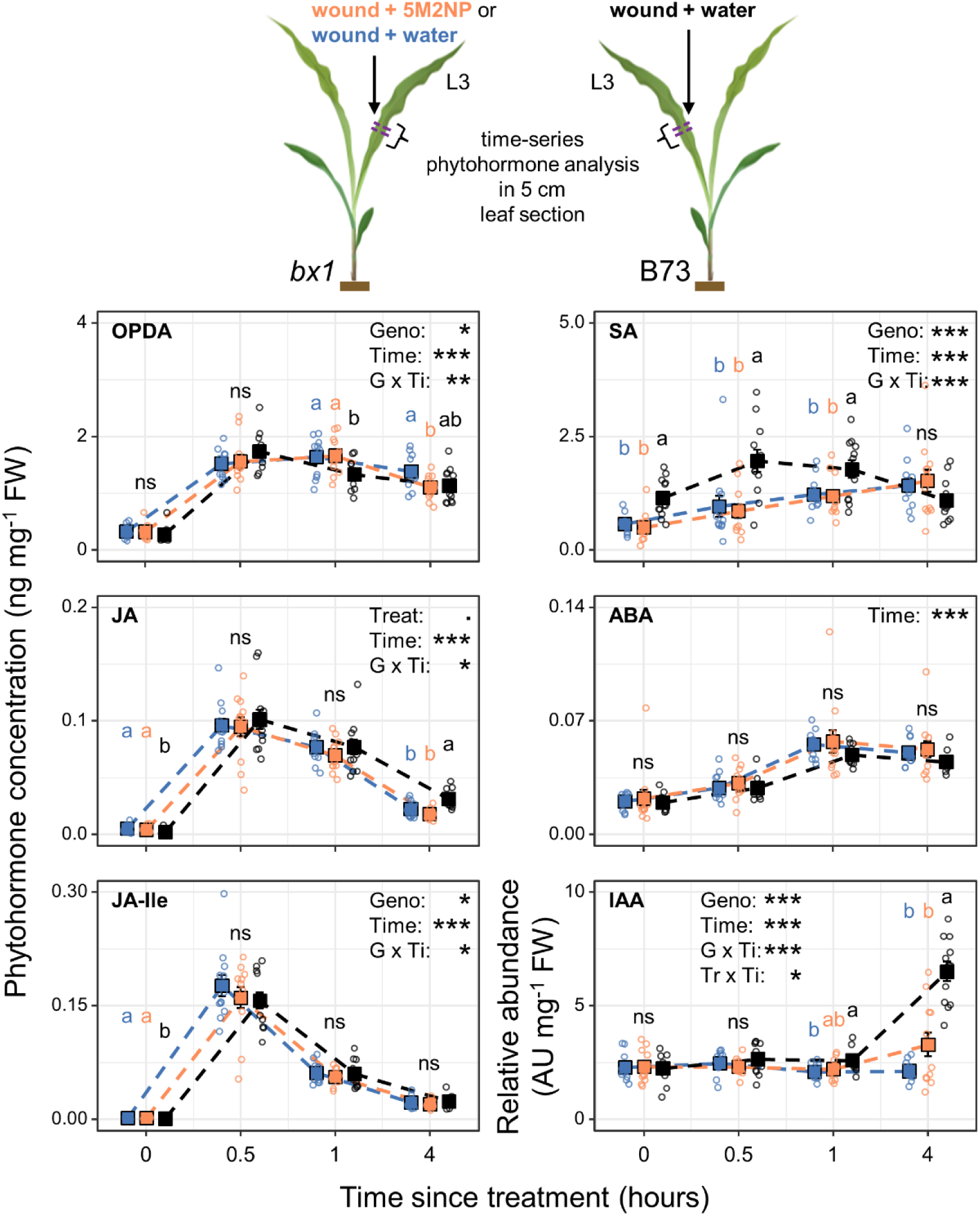
At a physiologically relevant dose, 5-Methoxy-2-Nitrophenol (5M2NP) has no major effect on concentrations of stress/defense-associated hormones in wounded leaf tissue. The figure shows local, time-resolved phytohormone accumulation profiles in leaves of maize (*Zea mays*) *bx1* mutants or the corresponding wild type, B73, following a wound + 5M2NP treatment (in orange) or a wound + water treatment (in blue for *bx1*, in black for B73). The third leaf (L3) of 16-d-old seedlings was wounded four times, perpendicular to but not crossing the midrib, using a hemostat. Wounds were immediately supplemented with 20 µl of a 5M2NP solution (7.5 ng 5M2NP µl^-1^) or Milli-Q water (0.75% v/v dichloromethane) as a control. At 0.5 h, 1 h and 4 h after these treatments, 5 cm long leaf sections centered around the wounds were cut out for phytohormone extraction. Corresponding leaf sections from untreated plants sampled at the beginning of the experiment (0 h) were used as controls. Squares depict mean (± SEM) concentrations of 12-oxo-phytodienoic acid (OPDA), jasmonic acid (JA), jasmonic acid–isoleucine (JA-Ile), salicylic acid (SA) and abscisic acid (ABA), as well as the relative abundance of indole-3-acetic acid (IAA), for each plant genotype and treatment, at indicated time points (n = 12). Black asterisks denote statistically significant effects of plant genotype (Geno or G), treatment (Tr), time since treatment (Time or Ti) and/or their interactions on the (relative) abundance of each hormone, as determined by a multifactorial analysis of variance (ANOVA): *** for *P* < 0.001; ** for *P* < 0.01; * for *P* < 0.05;. for *P* < 0.1; ns, not significant for *P* > 0.1. For clarity, only statistically significant effects are shown. Different letters indicate statistically significant differences between treatments and/or genotypes at individual time points, as determined by a one-way ANOVA, followed by estimated marginal means *post hoc* tests (Tukey-adjusted *P* ≤ 0.05). AU, arbitrary units; FW, fresh weight

To investigate if supplementation of wounds with 5M2NP enhances the emission of defense-associated volatiles, we repeated the experiment a third time and analyzed the plant’s headspace VOCs in real-time. As anticipated, mechanical wounding triggered a burst of GLVs, followed by the increased release of indole, various terpenes, and methyl salicylate (Fig. 4 and S9; factor Time). Moreover, supplementation of wounds with 5M2NP, as opposed to water, resulted in a stronger induction of monoterpene, sesquiterpene, homoterpene and methyl salicylate emissions (Fig. 4 and S9).

**Figure 4:**
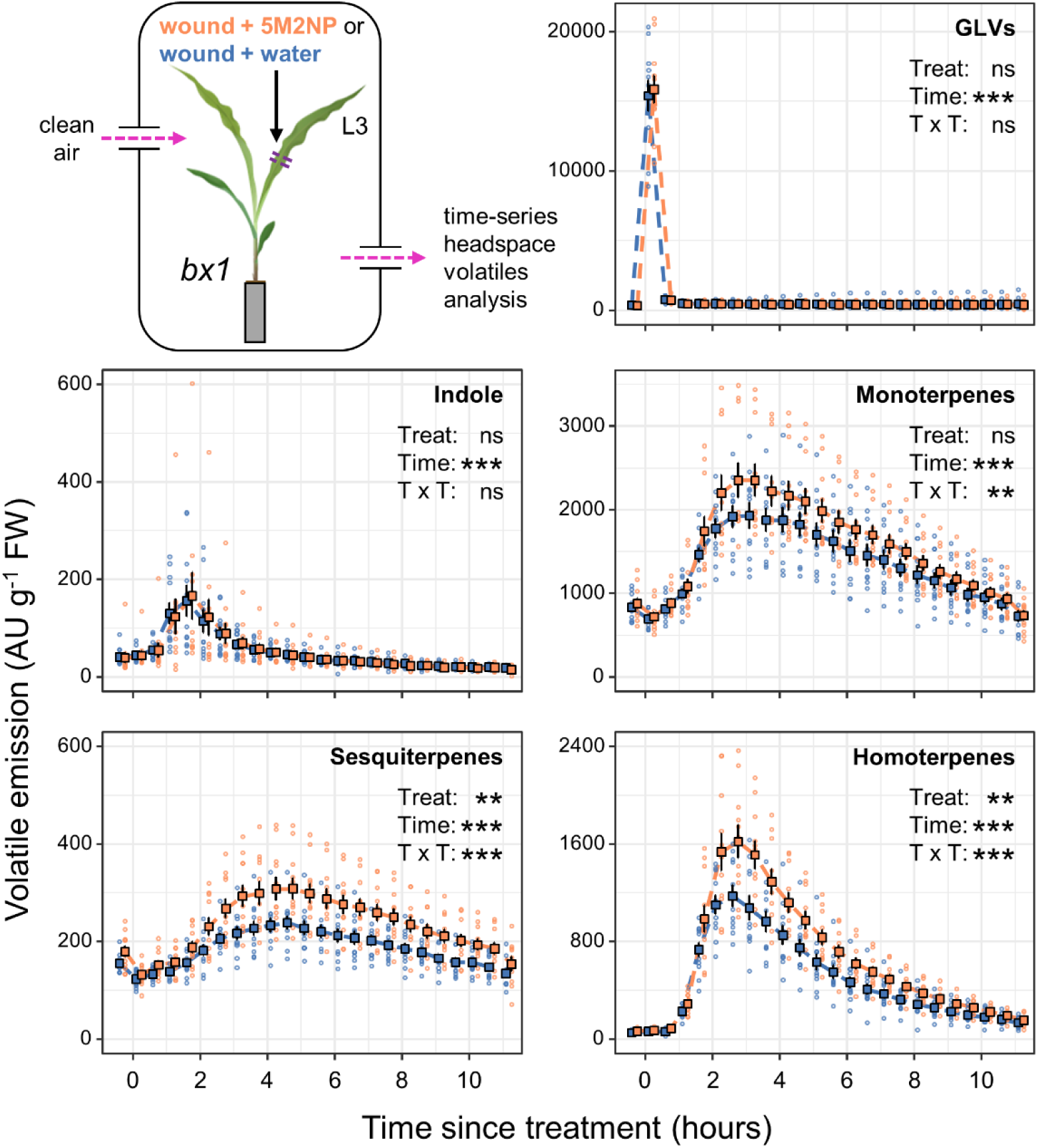
At a physiologically relevant dose, 5-Methoxy-2-Nitrophenol (5M2NP) enhances the wound-induced emission of defense-associated terpenoids. The figure shows time-resolved emission profiles of the main wound and defense-associated volatile organic compounds (VOCs) from maize (*Zea mays*) *bx1* mutants following either a leaf wound + 5M2NP treatment (in orange) or a wound + water treatment (in blue). The third leaf (L3) of 16-d-old *bx1* seedlings was wounded four times, perpendicular to but not crossing the midrib, using a hemostat. Wounds were immediately supplemented with 20 µl of a 5M2NP solution (7.5 ng 5M2NP µl^-1^) or Milli-Q water (0.75% v/v dichloromethane) as a control. Squares depict mean (± SEM) normalized emission values of green leaf volatiles (GLVs; sum of (*Z*)-3-hexenal, (*Z*)-3-hexen-1-ol, (*Z*)-3-hexenyl acetate), indole, monoterpenes, sesquiterpenes and homoterpenes (sum of DMNT and TMTT) for each treatment, at indicated time points (n = 11). Regarding the post-treatment data, black asterisks denote statistically significant effects of plant treatment (Treat), time since treatment (Time) and/or their interaction (T x T) on the emission of each VOC, or class of VOCs, as determined by a multifactorial repeated measures analysis of variance on aligned rank-transformed data (RM-ANOVA-ART): *** for *P* < 0.001; ** for *P* < 0.01; * for *P* < 0.05;. for *P* < 0.1.; ns, not significant for *P* > 0.1. Detailed results from the statistical analysis, including effects of plant treatment at individual time points, are listed in Table S2. Emission profiles of the individual GLVs and homoterpenes are depicted in Fig. S9. AU, arbitrary units; FW, fresh weight

Collectively, our results show that 5M2NP enhances defense gene expression and volatile release, but not phytohormone accumulation, suggesting that it acts as a defense modulator, either downstream from phytohormones or independently. We did not observe cell death in 5M2NP-supplemented wounds, suggesting 5M2NP is not strongly phytotoxic at the tested dose.

### 5M2NP acts as a direct defense against insect herbivores

To investigate whether 5M2NP directly affects the behavior and/or performance of arthropod herbivores, we conducted bioassays with four insect species, representing leaf or root-feeders with contrasting levels of adaptation to maize. When added to artificial diet (AD) at physiologically relevant doses, 5M2NP reduced the growth of caterpillar larvae of both generalist and specialist *Spodoptera* species (Lepidoptera: Noctuidae), but it did not deter them (Fig. 5). In choice essays, more *S. littoralis* larvae were located on/near food cubes supplemented with 1.0 ng 5M2NP mg^-1^ AD than on/near cubes supplemented with water (control). This preference for 5M2NP-treated AD was most pronounced after 960 min. At a higher concentration of 2.5 ng 5M2NP mg^-1^ AD, no clear preference was visible anymore (Fig. 5a). No clear preference was found for *S. frugiperda*, apart from the 960 min time point, when more larvae were found on/near diet cubes complemented with 1.0 ng 5M2NP mg^-1^ (Fig. 5c). In no-choice assays, caterpillars gained significantly less weight on 5M2NP-supplemented AD than on water-supplemented AD in a concentration-dependent manner. Compared to the water controls, the average weight of larvae exposed to 1.0 or 2.5 ng 5M2NP mg^-1^ AD for twelve consecutive days was reduced by 21 and 38%, respectively, for *S. littoralis* (Fig. 5b) and by 11 and 32%, respectively, for *S. frugiperda* (Fig. 5d).

**Figure 5:**
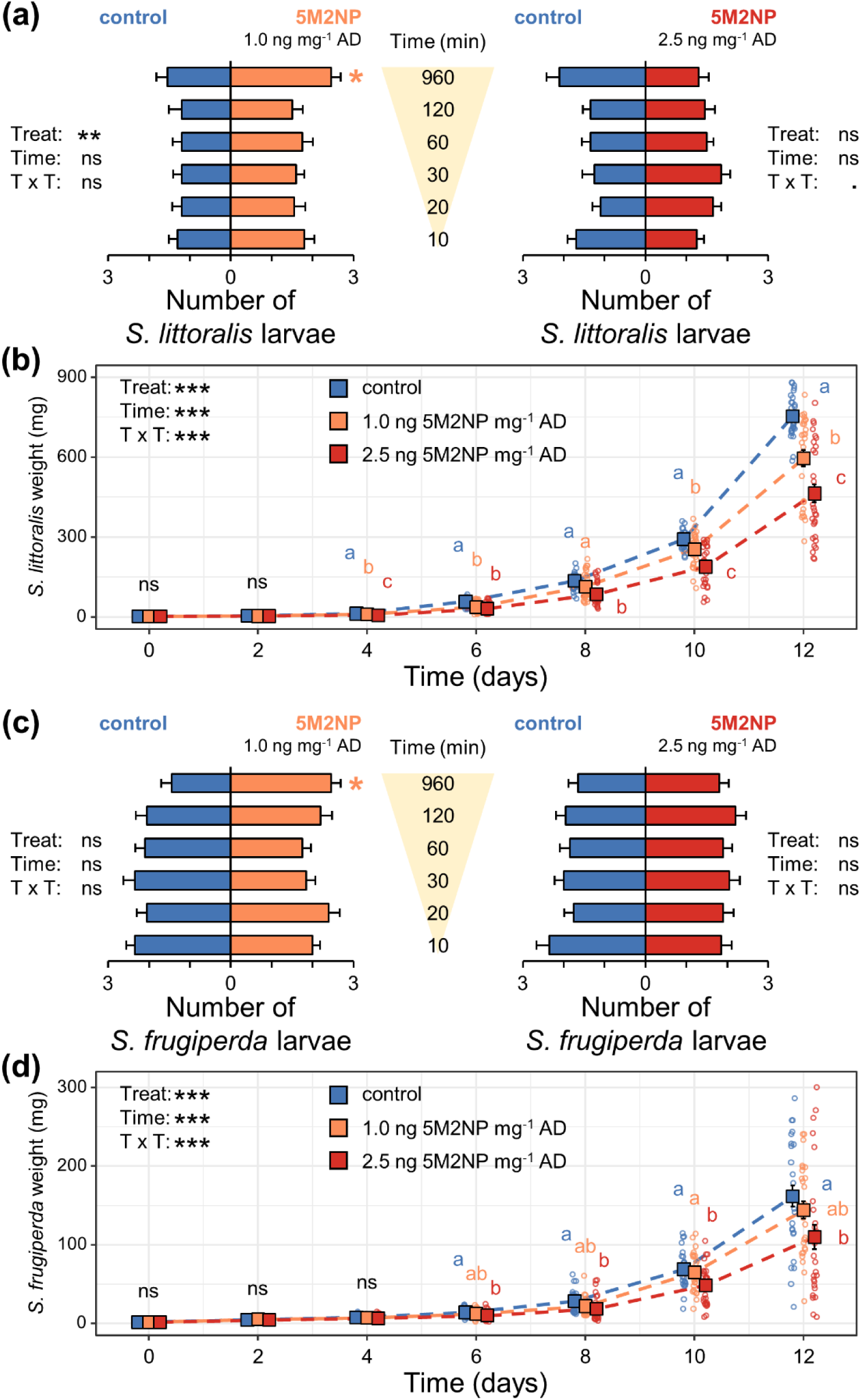
5-Methoxy-2-Nitrophenol (5M2NP) hampers the development of larvae of generalist and specialist *Spodoptera* sp., but does not deter them. When added to a soy-wheat germ-based artificial diet (AD) at physiological doses, 5M2NP alters the behavior and/or performance of naive (a, b) *Spodoptera littoralis* and (c, d) *Spodoptera frugiperda* larvae in a concentration-dependent manner. (a, c) Results from choice assays. Bars indicate mean (± SEM) number of larvae choosing AD supplemented with 5M2NP (at 1.0 or 2.5 ng mg^-1^ AD) versus AD supplemented with Milli-Q water (0.9% v/v ethanol) as a control, over time (n = 20 choice situations with five larvae each). (b, d) Results from no-choice assays. Squares indicate mean (± SEM) weight of larvae reared on AD supplemented with either 5M2NP (at 1.0 or 2.5 ng mg^-1^AD) or, as a control, on AD supplemented with Milli-Q water (0.9% v/v ethanol), over time (n = 30). Black asterisks denote statistically significant effects of AD treatment (Treat), time since the start of the bioassay (Time) and/or their interaction (T x T), as determined by (a, c) a Type II Wald chi-square test applied to a generalized linear mixed-effects model (GLMM) with Poisson distribution or (b, d) a multifactorial repeated measures analysis of variance on aligned rank transformed data (RM-ANOVA-ART): *** for *P* < 0.001; ** for *P* < 0.01; * for *P* < 0.05;. for *P* < 0.1.; ns, not significant for *P* > 0.1. Orange asterisks in panels (a, c) indicate significant differences between treatments at individual time points according to a one-sample *t*-test based on the percentage of larvae that chose for either of the AD treatments (* for *P* < 0.05). Different letters in panels (b, d) indicate statistically significant differences between treatments at individual time points, as determined by a one-way analysis of variance (ANOVA), followed by estimated marginal means *post hoc* tests (Tukey-adjusted *P* ≤ 0.05).

When added to the rhizosphere of *bx1* seedlings at a physiologically relevant dose, 5M2NP deterred rootworm larvae of both generalist and specialist *Diabrotica* species (Coleoptera: Chrysomelidae), whereas it did not reduce their growth (Fig. 6). In choice assays, the average number of *D. balteata* larvae that rejected a healthy *bx1* host was *c.* 2-fold higher upon supplementation of the rhizosphere with 25 ng 5M2NP (equivalent to *c*. 0.02 ng 5M2NP mg^-1^ root FW), as compared to supplementation with water (Fig. 6a). A similar trend was observed when 125 ng 5M2NP was added to the root system (equivalent to *c*. 0.11 ng 5M2NP mg^-1^ root FW; Fig. 6b). The repellent effect of 5M2NP on *D. v. virgifera* larvae was less strong and only evident for the lower of two doses tested, for which a *c.* 40% increase in the host-rejection rate was observed (Fig. 6d, e). The average host-rejection rate of control-treated plants was higher for *D. v. virgifera* than for *D. balteata* larvae, presumably because the former uses benzoxazinoid-containing metabolites as host-finding cues (Robert et al., 2012; Hu et al., 2018), which are largely absent from *bx1* roots. In no-choice assays, rootworm weight gain was similar on 5M2NP-supplemented and water-supplemented *bx1* seedlings (Fig. 6c, f). These bioassays indicate that 5M2NP has antibiotic and antixenotic properties towards chewing insect herbivores.

**Figure 6:**
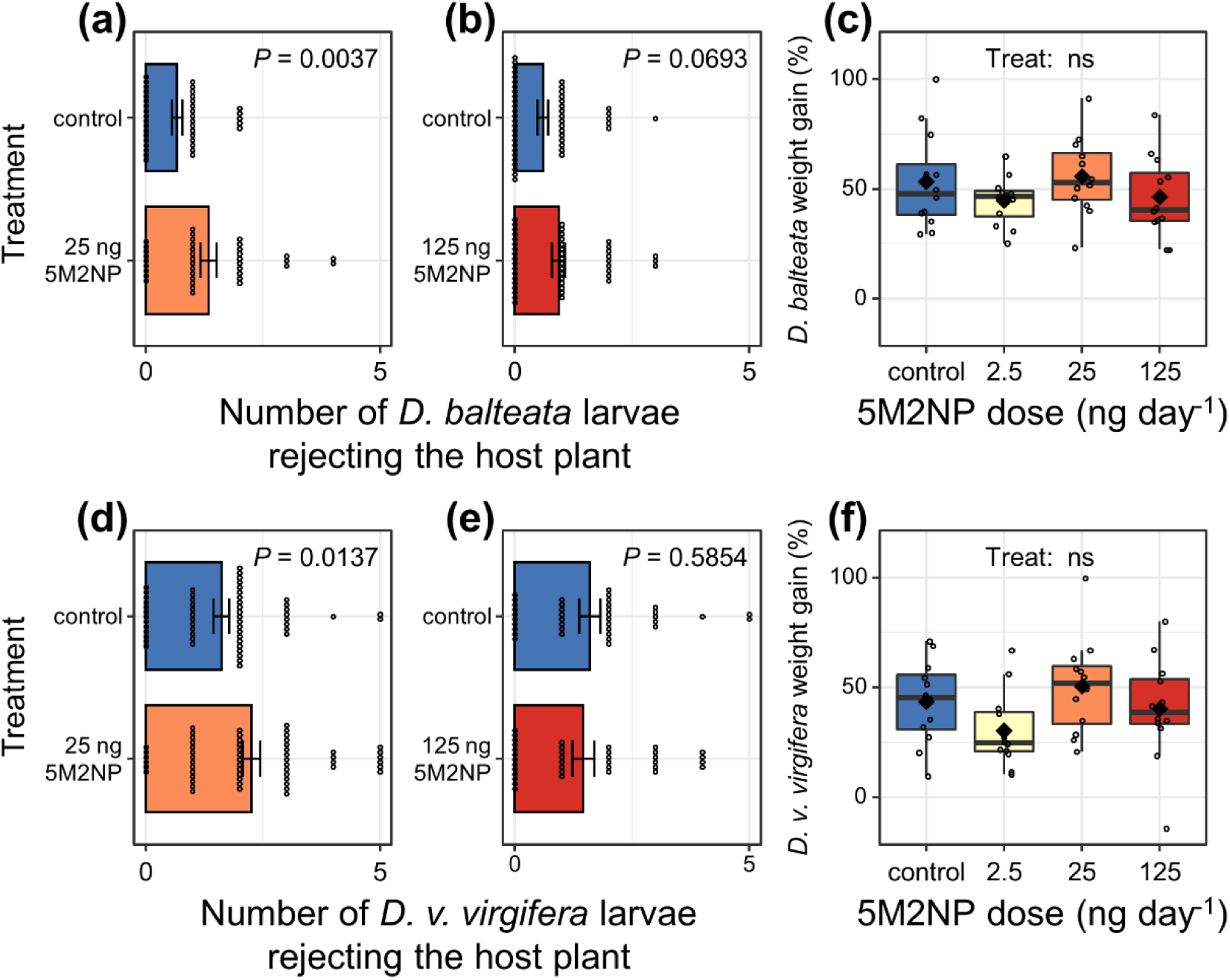
5-Methoxy-2-Nitrophenol (5M2NP) deters larvae of generalist and specialist *Diabrotica* sp., but does not hamper their development. When added to the root system of 16-d-old maize (*Zea mays*) *bx1* mutants at physiological doses, 5M2NP alters the behavior but not the performance of naive (a – c) *Diabrotica balteata* and (d – f) *Diabrotica virgifera virgifera* larvae in a concentration-dependent manner. (a, b, d, e) Results from choice assays. Bars indicate mean (± SEM) number of larvae rejecting *bx1* seedlings, rhizo-supplemented with either 5M2NP (25 ng or 125 ng) or with autoclaved tap water (0.0025% v/v ethanol) as a control (n = 37 – 58 choice situations with five larvae each). *P*-values represent the outcome of a Type II Wald chi-square test applied to a generalized linear mixed-effects model (GLMM) with Poisson distribution and reflect the significance of plant treatment on *Diabrotica* larval behavior. Note that most data points from the control treatments in panels (a) and (b) are identical, i.e., shared, see Methods S5 for details. The same applies to data in panels (d) and (e). (c, f) Results from no-choice assays. Boxplots show percent weight gain, over the course of 4 d, of larvae reared on *bx1* seedlings, rhizo-supplemented with either 5M2NP (2.5 ng, 25 ng or 125 ng day^-1^) or with autoclaved tap water (0.0025% v/v ethanol) as a control (n = 12 plants with five larvae each). Boxes span the 25 – 75 percentiles, whiskers span the 1.5x interquartile range. Horizontal lines and diamonds inside the boxes represent medians and means, respectively. The effect of the 5M2NP treatment (Treat) on larval weight gain was not statistically significant (ns; *P* > 0.1), as determined by a one-way analysis of variance (ANOVA).

## DISCUSSION

Plants produce a plethora of bioactive secondary metabolites (Dixon, 2001; Mithöfer and Boland, 2012; Osbourn and Lanzotti, 2009). Tissue disruption can trigger the degradation of these metabolites, and the resulting catabolites further increase the plant’s chemical and functional repertoire. Yet, most breakdown products of plant secondary metabolites remain unknown, despite their potentially significant biological activities. Here we identified 5-Methoxy-2-Nitrophenol (5M2NP), as a benzoxazinoid breakdown product that accumulates in damaged maize tissues and acts both as a direct defense and as a defense regulator. Methoxy-nitrophenols have not been described before in plants, and our work therefore expands our understanding of the chemical diversity of plant natural products as well as the chemical biology of plant defense.

Upon tissue disruption, plant secondary metabolites such as glucosinolates, cyanogenic glycosides, sesquiterpene lactone glycosides, triterpene saponins, and benzoxazinoids are converted into biologically active compounds (Wouters et al., 2016a; Halkier and Gershenzon, 2006; Mithöfer and Boland, 2012; Lacchini et al., 2023). In the case of benzoxazinoids, wounding triggers the conversion of DIMBOA-Glc into the reactive DIMBOA, which further decays into MBOA (Wouters et al., 2016b). However, MBOA accumulation does not account for the entirety of DIMBOA disappearance (Woodward et al., 1978), suggesting that other products may be formed. Here, we identify 5M2NP as an additional DIMBOA catabolite, and thus add a novel branch to the benzoxazinoid degradation pathway.

How is 5M2NP formed? Our experiments show that DIMBOA can be converted into 5M2NP (Fig. 1), both in the presence and absence of plant tissue, but the conversion rate of 5M2NP is much higher in the presence of plant material. The latter can be explained by the presence and activity of one or more hitherto unknown enzymes in plant material that catalyze this conversion. Alternatively or additionally, environmental parameters such as changes in pH or solubility may accelerate 5M2NP formation in disrupted plant cells (Wouters et al., 2016b). Although additional research is needed to determine how and when 5M2NP accumulates *in planta*, we speculate that DIMBOA’s oxo-cyclo ring-chain tautomerism, i.e. the continuous interconversion of DIMBOA with a closed versus open ring structure, is critical for the formation of 5M2NP. That is, breakage of the nitrogen-carbonyl bond in open-form DIMBOA and subsequent oxidation of the putative 5-Methoxy-2-Hydroxyaminophenol intermediate will likely yield 5M2NP (Fig. S10). The chemical transformation in the reverse direction is well established, as synthetic 5M2NP is commonly used as substrate to produce benzoxazinoids, including DIMBOA (Jernow and Rosen, 1973; Campos et al., 1989). However, this reverse reaction is unlikely to occur *in planta*.

Breakdown products of secondary metabolites can act as regulators of defense responses (Erb and Kliebenstein, 2020). Despite the low abundance of 5M2NP in damaged tissues, our experiments demonstrate its potency as a defense modulator. In micromolar quantities, exogenously applied 5M2NP boosted the local wound-induced expression of maize genes, including genes coding for defensive proteinase inhibitors and enzymes involved in the biosynthesis of jasmonates and volatile terpenoids (Fig. 2). Suppressed transcript levels of benzoxazinoid biosynthesis genes *Bx10* and *Bx11* suggest that 5M2NP may also be involved in retrograde signaling, as has been proposed for DIMBOA (Ahmad et al., 2011). We did not detect an effect of 5M2NP on the accumulation of jasmonates in damaged leaf sections (Fig. 3). Since our first sampling point was 30 min after treatments – coinciding with the peak of the wound-induced JA burst (Engelberth et al., 2007) – it is conceivable that we have missed an earlier 5M2NP-mediated effect on JA concentrations; for example, a faster production, reminiscent of a priming response (Martinez-Medina et al., 2016). Alternatively, 5M2NP might modulate defenses independently-or downstream from jasmonates. Either way, 5M2NP amplified the wound-induced emission of volatile mono, homo and sesquiterpenes (Fig. 4), several of which are JA-inducible (Li et al., 2015b; Lenk et al., 2012) and involved in defense regulation and/or indirect defense (Vlot et al., 2021; Brosset and Blande, 2022; Turlings and Erb, 2018). Like with the gene expression, 5M2NP influenced some but not all monitored headspace VOCs, indicating specific regulation of a subset of defense responses.

Multifunctionality is increasingly viewed as a common property of plant secondary metabolites (Neilson et al., 2013; Piasecka et al., 2015; Erb and Kliebenstein, 2020; Weng et al., 2021). For example, volatile indole – produced and emitted upon herbivory (Turlings et al., 1990; Frey et al., 2000) – deters young *S. littoralis* larvae and increases their mortality (Veyrat et al., 2016), and simultaneously primes the accumulation of JA-Ile, ABA and volatile terpenes (Erb et al., 2015). Here, we find that 5M2NP acts both as a defense modulator and as a direct defense. In this regard, the capacity of 5M2NP to suppress caterpillar growth (Fig. 5) and deter rootworms (Fig. 6) at micro and nanomolar quantities, respectively, deserves attention. As hydrolysis product of the most abundant benzoxazinoid in undamaged maize leaves of many maize cultivars (Glauser et al., 2011; Meihls et al., 2013), DIMBOA suppresses growth of *S*. *littoralis* at concentrations of 200 µg DIMBOA g^-1^ artificial diet, i.e. 2.0 x 10^8^ ng DIMBOA mg^-1^ artificial diet (Glauser et al., 2011). Yet, even at this high concentration, maize-adapted *S. frugiperda* is not affected by DIMBOA (Glauser et al., 2011), presumably because it is able to rapidly re-glycosylate any consumed DIMBOA (Wouters et al., 2014). To what extent the re-glycosylation of DIMBOA by *S. frugiperda* prevents the *in vivo* formation of 5M2NP remains to be determined, but because *S. frugiperda* can develop equally well on WT and *bx1* mutant plants (Hu et al., 2021), it seems capable of avoiding negative effects of 5M2NP. Below ground, the western corn rootworm is a similarly specialized maize feeder who benefits from the presence of benzoxazinoids (Robert et al., 2012; Machado et al., 2021; Hu et al., 2018) and can guide their metabolization in its gut (Robert et al., 2017). Since *D. v. virgifera* larvae can transform highly reactive benzoxazinoids such as HMBOA (Robert et al., 2017), they may be able to avoid 5M2NP accumulation in their gut. Given 5M2NP’s deterrent effect on rootworms, it is tempting to speculate that 5M2NP can also function as indirect defense by serving as a host-finding cue for natural enemies of herbivores. However, preliminary experiments with entomopathogenic nematodes, which prey on rootworms, did not provide evidence to support this hypothesis (Fig. S11).

What is 5M2NP’s mode of action? Unlike its precursor DIMBOA, 5M2NP is not a reactive electrophile species, or at least not a potent one. We therefore expect that the nitro group is instrumental for 5M2NP’s functions, as nitroarenes are known to engage in redox processes that are responsible for mutagenic and/or genotoxic effects (Nepali et al., 2019; Tokiwa and Ohnishi, 1986). Analogously, the non-volatile 9-LOX-derived oxylipins 10-oxo-11-phytoenoic acid (10-OPEA) and 9-hydroxy-10-oxo-12(*Z*),15(*Z*)-octadecadienoic acid (9,10-KODA) accumulate in wounded maize tissues (Christensen et al., 2015; He et al., 2020; Yuan et al., 2023) and both metabolites promote defense signaling and are directly toxic to insect herbivores (Yuan et al., 2023; Christensen et al., 2015). The 13-LOX-derived GLVs are similarly multifunctional (Scala et al., 2013). Again, while 10-OPEA and some GLVs are reactive electrophile species, this cannot explain all functional properties of these and other oxylipins (Farmer and Mueller, 2013; Christensen et al., 2015; Scala et al., 2013). With regard to transcriptional regulation, DAMPs commonly signal through cell-surface-localized receptors (Tanaka and Heil, 2021), which allows for specific downstream responses. Yet, small molecules, such as 5M2NP, may also interact (in)directly with transcription factors, chromatin-modifying proteins, or enzymes to modulate their activity and, hence, gene expression (Venturelli et al., 2015; Li et al., 2023). Establishing the molecular mechanisms underlying 5M2NP’s toxicity and function as a defense regulator is an exciting prospect of this work. More generally, knowledge of the full functional repertoire of plant secondary metabolites will help to understand their ecology and evolution and lead to better applications for sustainable agriculture (Shen et al., 2023; Weng et al., 2021).

In conclusion, our work identifies a methoxy-nitrophenol as a potent direct defense and as a positive regulator of maize defense responses. The molecule is derived from benzoxazinoid catabolism and may thus accumulate at wound sites and in the digestive tracts of herbivores, especially of non-adapted ones, to unfold its actions. This discovery improves our understanding of the chemical defense arsenal of plants and adds another potentially important facet to the multiple beneficial functions of benzoxazinoids for cereals.

## DATA AVAILABILITY

All raw data related to this manuscript have been uploaded to XXXX and are publicly available. R code for statistical analysis and data visualization is available at GitHub: XXXX

## Supporting information

Schimmel_et_al_Supporting_Information

## ABBREVIATIONS

DIMBOA: 2,4-Dihydroxy-7-Methoxy-2H-1,4-Benzoxazin-3(*4H*)-One
DIM2BOA-Glc: (*2R*)-β-D-Glucopyranosyl-2,4-Dihydroxy-7,8-Dimethoxy-2H-1,4-Benzoxazin-3(*4H*)-One
DIMBOA-Glc: (*2R*)-β-D-Glucopyranosyl-2,4-Dihydroxy-7-Methoxy-2H-1,4-Benzoxazin-3(*4H*)-One
DMNT: (*3E*)-4,8-Dimethyl-1,3,7-Nonatriene
HDM2BOA-Glc: (*2R*)-β-D-Glucopyranosyl-2-Hydroxy-4,7,8-Trimethoxy-2H-1,4-Benzoxazin-3(*4H*)-One
HDMBOA-Glc: (*2R*)-β-D-Glucopyranosyl-2-Hydroxy-4,7-Dimethoxy-2H-1,4-Benzoxazin-3(*4H*)-One
HM2BOA-Glc: (*2R*)-β-D-Glucopyranosyl-2-Hydroxy-7,8-Dimethoxy-2H-1,4-Benzoxazin-3(*4H*)-One
MBOA: 6-Methoxy-Benzoxazolin-2(*3H*)-One
TMTT: (*3E,7E*)-4,8,12-Trimethyl-1,3,7,11-Tridecatetraene

## ACKNOWLEDGEMENTS

This work was financially supported by the European Research Council (ERC) under the European Union’s Horizon 2020 research and innovation programme (Marie Skłodowska-Curie grant agreement #794947 to B.C.J.S. and #886651 to L.W.; ERC-2019-STG949595 to C.A.M.R.), the Swiss National Science Foundation (#310030_189071 to C.A.M.R.; #192564 to M.E.), and the University of Bern. The authors wish to thank Tobias Züst and Mirco Hecht for technical assistance, and Christopher Ball and Jasmin Sekulovski for taking care of plants.

## AUTHOR CONTRIBUTIONS

B.C.J.S and M.E. conceived and designed experiments; B.C.J.S., R.E.B., C.C., V.O., L.W., A.V.M. and M.E. conducted experimental work; P.M. and C.A.M.R. contributed essential materials for experiments; B.C.J.S., R.E.B., P.M., G.G., L.W., C.A.M.R. and M.E. collected, analyzed and/or interpreted data. B.C.J.S. visualized the data. B.C.J.S. and M.E. wrote the manuscript with input from all the other authors.

## SUPPORTING INFORMATION

**Figure S1:** Identification of 5M2NP by means of SPME-GC-MS

**Figure S2:** 5M2NP (relative) abundance in leaves and roots of herbivore-attacked and non-attacked control maize seedlings

**Figure S3:** Identification and quantification of 5M2NP by means of UHPLC-MS/MS

**Figure S4:** Maize root 5M2NP content of seedlings grown from surface-sterilized seeds in soil-free conditions or grown in sand and soil

**Figure S5:** 5M2NP concentration in leaves of maize *bx1* mutants

**Figure S6:** 5M2NP accumulates in ground root tissue of *bx1* mutants supplemented with DIMBOA

**Figure S7:** Relative abundance of 5M2NP in roots of seedlings subjected to pharmacological treatments aimed at modifying the metabolic flux through the phenylpropanoid pathway

**Figure S8:** Concentrations of 5M2NP and eleven benzoxazinoids in the crown roots of *brown-midrib* mutants (*bm1-4*) and their corresponding wild type F2

**Figure S9:** 5M2NP enhances the wound-induced emission of maize VOCs

**Figure S10:** Hypothetical metabolic route from DIMBOA to 5M2NP

**Figure S11:** 5M2NP does not alter the foraging behavior of *Heterorhabditis bacteriophora* entomopathogenic nematodes (EPNs) in Petri dish-based choice assays

**Table S1:** qRT-PCR primer specifications

**Table S2:** Results from statistical analysis of maize headspace VOCs data, presented in Figs. 4 and S9.

**Methods S1:** 5M2NP quantification by means of UHPLC-MS/MS

**Methods S2:** Gene expression analysis

**Methods S3:** Phytohormone analysis

**Methods S4:** Caterpillar preference assays

**Methods S5:** Rootworm preference assays

**Methods S6:** Caterpillar performance assays

**Methods S7:** Rootworm performance assays

**Methods S8:** Plant infestations with herbivores

**Methods S9**: EPN preference assays

## CITATION DIVERSITY STATEMENT

Recent work in several fields of science, including plant science, has identified a consistent bias in citation practices such that papers from women and other minority scholars are under-cited relative to the number of such papers in the field (Mitchell et al., 2013; Marks et al., 2023; Caplar et al., 2017; Dion et al., 2018; Dworkin et al., 2020; Wang et al., 2021; Chatterjee and Werner, 2021). Here, inspired by a growing social movement to overcome this bias (Lloreda, 2022; Kwon, 2022), we sought to proactively consider choosing references that reflect the diversity of the field in thought, form of contribution, gender, race, ethnicity, and other factors. Additionally, we sought to raise awareness about the citation bias by reporting on (presumed) gender and ethnicity relations of the literature cited in this work (Zurn et al., 2020; Zurn et al., 2022). To do so, we obtained the predicted gender of the first and last author of each reference by using databases that store the probability of a first name being carried by a woman (Dworkin et al., 2020; Zhou et al., 2020). By this measure – and excluding self-citations to the first and last authors of this manuscript as well as citations used in this statement – our references contain 5% woman(first)/woman(last), 10% man/woman, 37% woman/man, and 48% man/man. This method is limited in that a) names, pronouns, and social media profiles used to construct the databases may not, in every case, be indicative of gender identity, b) it cannot account for intersex, non-binary, or transgender people, and c) shared first-or senior authorships are not taken into account. Second, we obtained predicted racial/ethnic category of the first and last author of each reference by databases that store the probability of a first and last name being carried by an author of color (Ambekar et al., 2009; Chintalapati et al., 2023). By this measure (and excluding self-citations), our references contain 13% author of color (first)/author of color(last), 9% white author/author of color, 24% author of color/white author, and 54% white author/white author. This method is limited in that a) names and Florida Voter Data to make the predictions may not be indicative of racial/ethnic identity, b) it cannot account for Indigenous and mixed-race authors, or those who may face differential biases due to the ambiguous racialization or ethnicization of their names, and and c) shared first-or senior authorships are not taken into account.. We look forward to future work that could help us to better understand how to support equitable practices in science. Instructions, code and a template for generating a Citation Diversity Statement can be found at: https://github.com/dalejn/cleanBib

