## Supplementary material for "The benzoxazinoid breakdown product 5-Methoxy-2-Nitrophenol has a dual function in maize defense against herbivory": Schimmel_et_al_Supporting_Information

**TITLE**

Bernardus C. J. Schimmel

Matthias Erb

**CONTENTS**

**Figure S1:** Identification of 5M2NP by means of SPME-GC-MS
**Figure S2:** 5M2NP (relative) abundance in leaves and roots of herbivore-attacked and non-attacked control maize seedlings
**Figure S3:** Identification and quantification of 5M2NP by means of UHPLC-MS/MS
**Figure S4:** Maize root 5M2NP content of seedlings grown from surface-sterilized seeds in soil-free conditions or grown in sand and soil
**Figure S5:** 5M2NP concentration in leaves of maize *bx1* mutants
**Figure S6:** 5M2NP accumulates in ground root tissue of *bx1* mutants supplemented with DIMBOA
**Figure S7:** Relative abundance of 5M2NP in roots of seedlings subjected to pharmacological treatments aimed at modifying the metabolic flux through the phenylpropanoid pathway
**Figure S8:** Concentrations of 5M2NP and eleven benzoxazinoids in the crown roots of *brown-midrib* mutants (*bm1-4*) and their corresponding wild type F2
**Figure S9:** 5M2NP enhances the wound-induced emission of maize VOCs
**Figure S10:** Hypothetical metabolic route from DIMBOA to 5M2NP
**Figure S11:** 5M2NP does not alter the foraging behavior of *Heterorhabditis bacteriophora* entomopathogenic nematodes (EPNs) in Petri dish-based choice assays

**Table S1:** qRT-PCR primer specifications
**Table S2:** Results from statistical analysis of maize headspace VOCs data, presented in Figs. 4 and S9.

**Methods S1:** 5M2NP quantification by means of UHPLC-MS/MS
**Methods S2:** Gene expression analysis
**Methods S3:** Phytohormone analysis
**Methods S4:** Caterpillar preference assays
**Methods S5:** Rootworm preference assays
**Methods S6:** Caterpillar performance assays
**Methods S7:** Rootworm performance assays
**Methods S8:** Plant infestations with herbivores
**Methods S9**: EPN preference assays

(a)

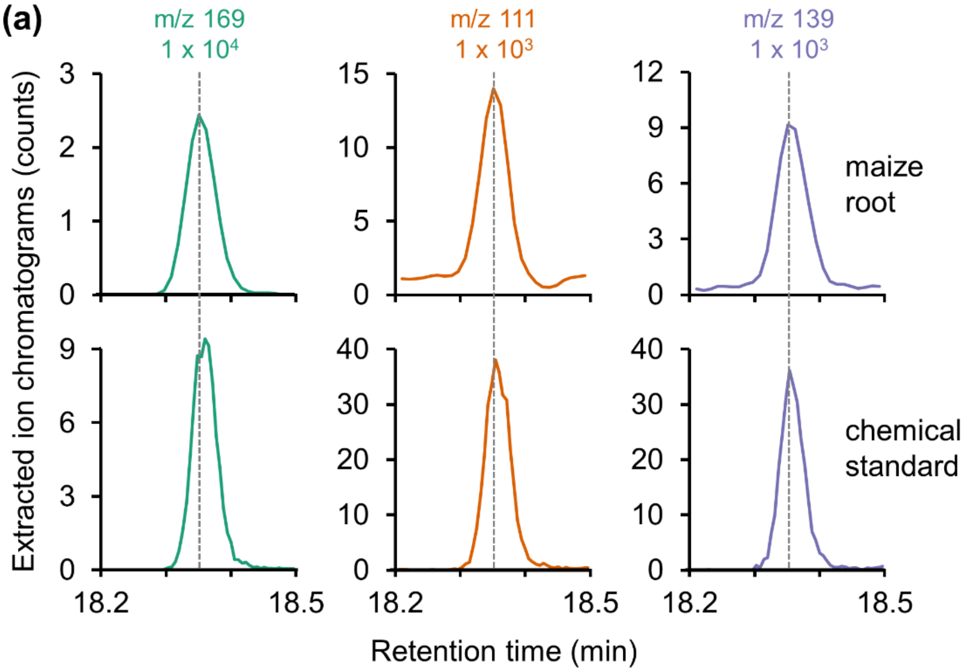
m/z 169
1 x 10⁴
m/z 111
1 x 10³
m/z 139
1 x 10³
maize root
chemical standard
Extracted ion chromatograms (counts)
18.2
18.5
Retention time (min)

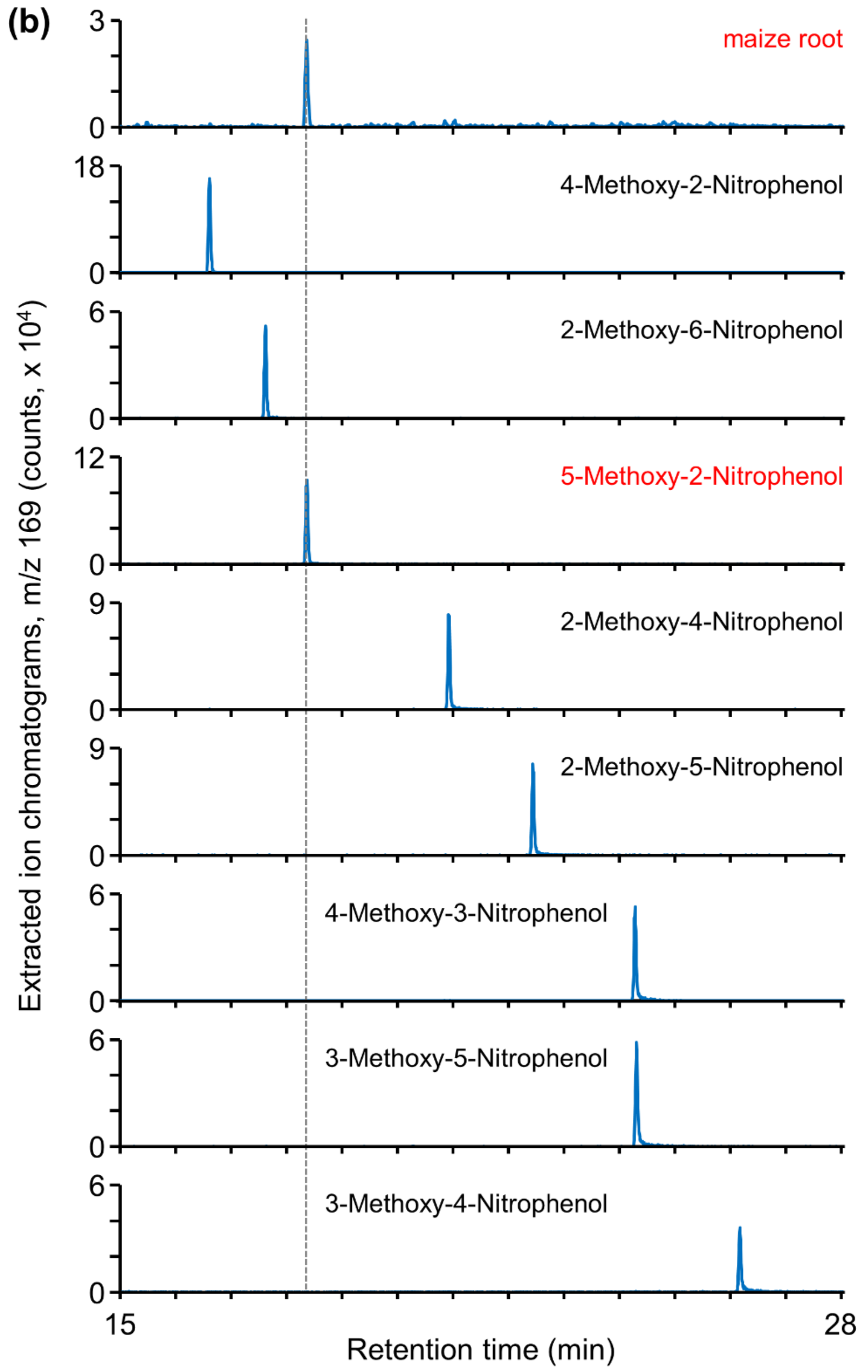


**Figure S1:** Identification of 5-Methoxy-2-Nitrophenol (5M2NP) in ground maize (*Zea mays*) roots by means of SPME-GC-MS. (a) Representative GC-MS extracted ion chromatograms showing diagnostic electron ionization ions of 5M2NP (*m/z* 169, *m/z* 111 and *m/z* 139) as detected in volatiles emanating from *c*. 100 mg ground crown roots of 21-d-old maize plants (cv. Delprim; top panels) and from 2 ng of 5M2NP chemical standard (bottom panels). (b) GC-MS extracted ion chromatograms of the main diagnostic electron ionization ion for methoxy-nitrophenols (*m/z* 169) and relative retention times of volatiles emanating from approximately 100 mg ground crown roots of 21-d-old maize plants (cv. Delprim) and from various methoxy-nitrophenol chemical standards (2 ng each).

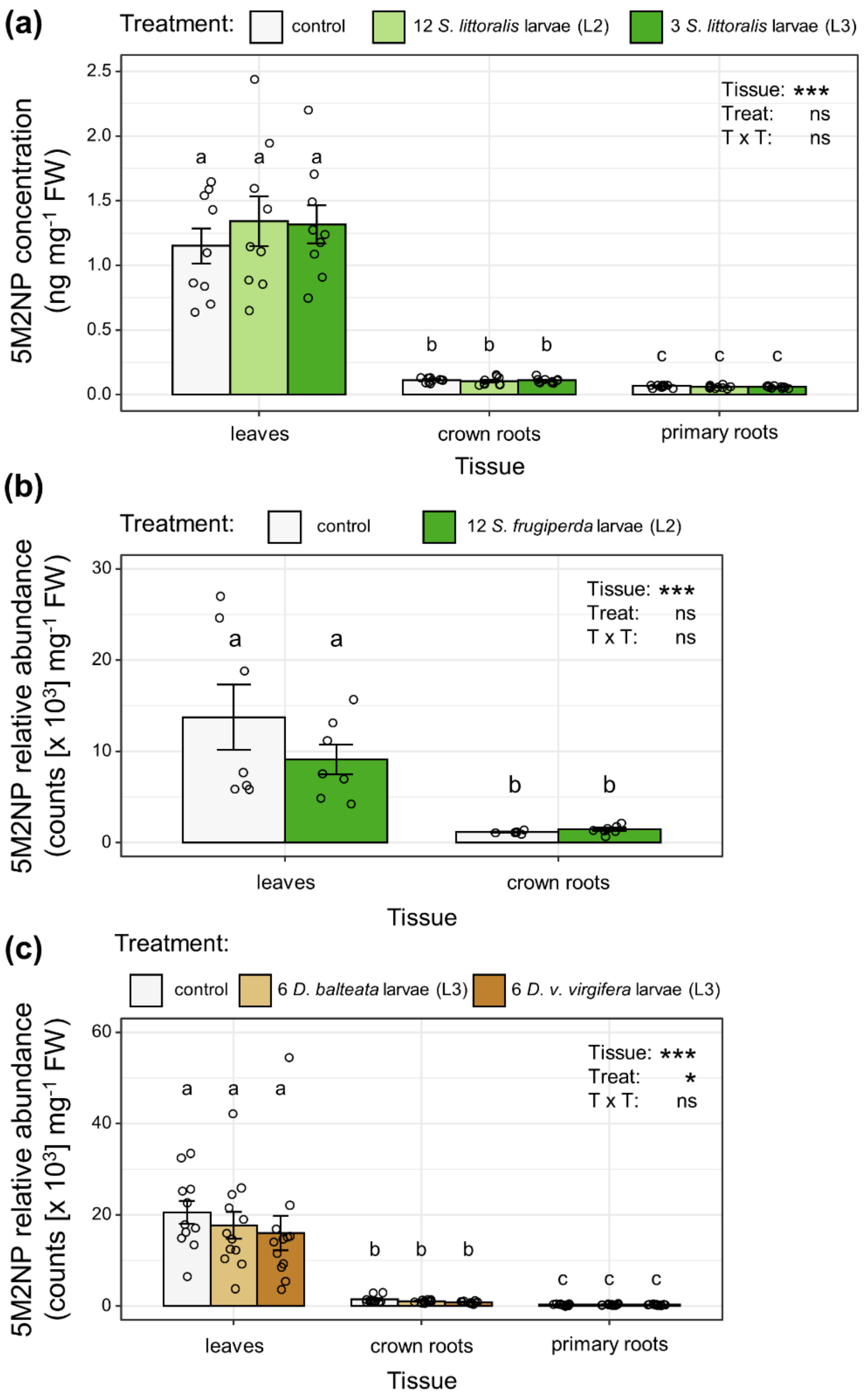


**Figure S2:** The (relative) abundance of 5-Methoxy-2-Nitrophenol (5M2NP) in leaves and roots of herbivore-attacked and non-attacked control maize (*Zea mays*) seedlings. Bars indicate mean (± SEM) amounts of 5M2NP in designated tissues of: (a) 23-d-old Delprim plants infested aboveground with second or third-instar *Spodoptera littoralis* larvae for 2 d (n = 9 – 10); (b) 16-d-old B73 plants infested aboveground with second-instar *Spodoptera frugiperda* larvae for 2 d (n = 7); and (c) 16-d-old Delprim plants infested belowground with third-instar *Diabrotica balteata* or *Diabrotica virgifera virgifera* larvae for 4 d (n = 12). Corresponding tissues from healthy non-infested plants were harvested as controls. Black asterisks denote statistically significant effects of plant tissue (Tissue), treatment (Treat) and/or their interaction (T x T) on the 5M2NP abundance, as determined by a two-way analysis of variance (ANOVA; panels a, b) or by a Type II Wald chi-square test applied to a linear mixed model (LMM; panel c): *** for $P < 0.001$; ** for $P < 0.01$; * for $P < 0.05$; . for $P < 0.1$.; ns, not significant for $P > 0.1$. Different letters above bars indicate statistically significant differences between plant tissues and/or treatments as determined by estimated marginal means *post hoc* tests (Tukey-adjusted $P \leq 0.05$). See Methods S8 for experimental details about plant infestations with herbivores. FW, fresh weight

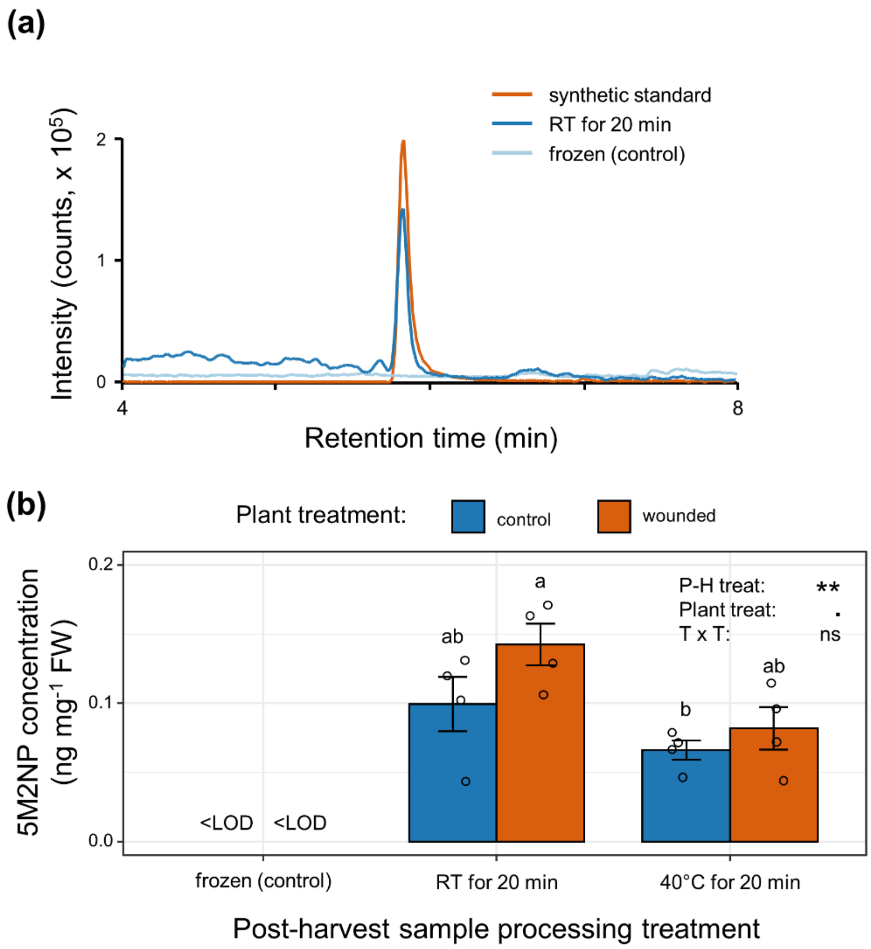


**Figure S3:** Identification and quantification of 5-Methoxy-2-Nitrophenol (5M2NP) in ground maize (*Zea mays*) leaves by means of UHPLC-MS/MS. (a) Representative HPLC-MS/MS chromatograms of a diagnostic transition of 5M2NP (*m/z* 168>153) as detected in methanol extracts of *c*. 100 mg ground homogenized leaf tissue of 14-d-old B73 seedlings and from a synthetic standard (50 ng 5M2NP $ml^{-1}$). (b) 5M2NP concentration in leaves of mechanically wounded or undamaged control leaves of 14-d-old B73 seedlings. Bars indicate mean (± SEM) amounts of 5M2NP (n = 4). Prior to 5M2NP extraction, sample aliquots, containing *c*. 100 mg of ground frozen homogenized leaf material produced from each biological replicate, were subjected to a secondary treatment. That is, one aliquot was incubated at room temperature (RT) for 20 min, thus allowing the tissue to thaw. A second aliquot was incubated at 40°C for 20 min, while a third aliquot was kept at -80°C as a control. Black asterisks denote statistically significant effects of plant treatment (Plant treat), post-harvest treatment (P-H treat) and/or their interaction (T x T) on the 5M2NP concentration, as determined by a two-way analysis of variance (ANOVA): *** for $P < 0.001$; ** for $P < 0.01$; * for $P < 0.05$; . for $P < 0.1$.; ns, not significant for $P > 0.1$. Different letters above bars indicate statistically significant differences between treatments as determined by estimated marginal means *post hoc* tests (Tukey-adjusted $P \leq 0.05$). The limit of detection (LOD) using the applied UHPLC-MS/MS methodology was 3 ng 5M2NP $ml^{-1}$ or 0.015 ng 5M2NP $mg^{-1}$ fresh weight (FW).

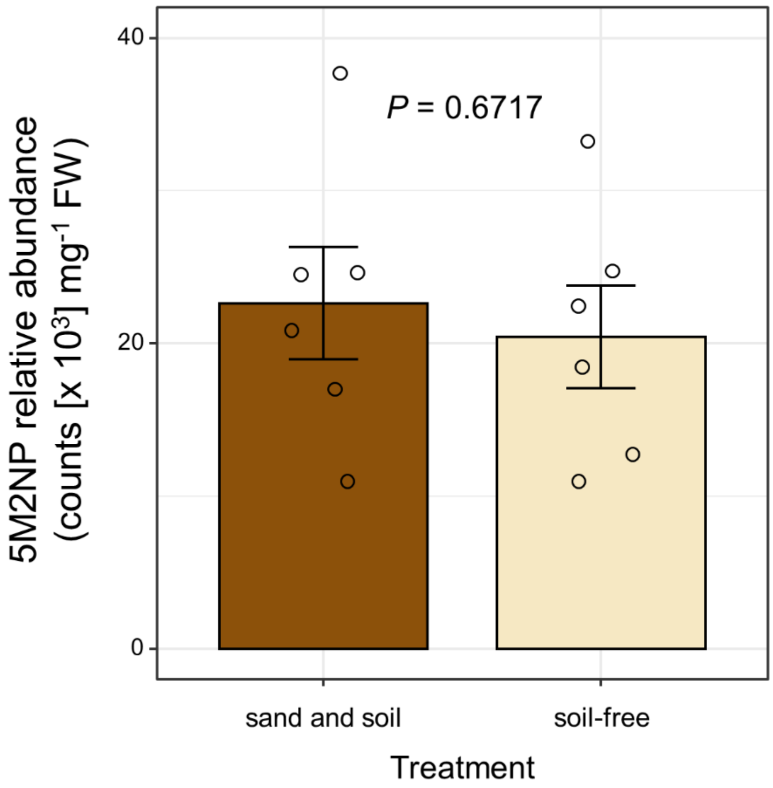


**Figure S4:** The root 5-Methoxy-2-Nitrophenol (5M2NP) content of maize (*Zea mays*) seedlings grown from surface-sterilized seeds in soil-free conditions is similar to that of maize seedlings grown in sand and soil (i.e. standard conditions). Bars indicate mean (± SEM) amounts of 5M2NP in crown roots of 16-d-old W22 plants subjected to the distinct growth conditions (n = 6). The *P*-value represents the outcome of a two-sided Student's *t-test*. FW, fresh weight

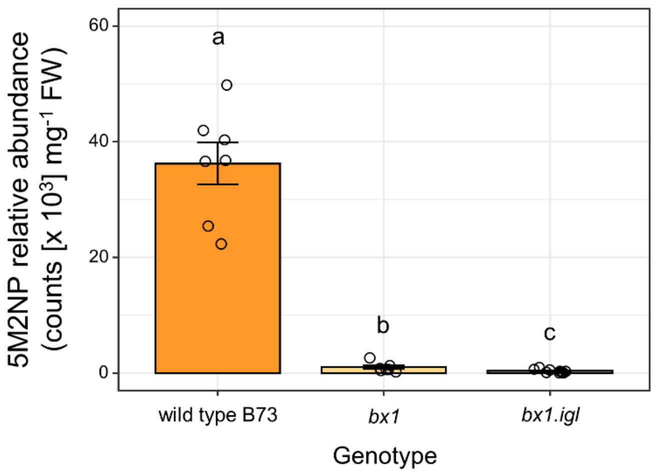


**Figure S5:** Compared to wild type maize (*Zea mays*), the 5-Methoxy-2-Nitrophenol (5M2NP) concentration is greatly reduced in leaves of *bx1* mutants. Bars indicate mean (± SEM) amounts of 5M2NP in leaves of 16-d-old *bx1* single and double mutants along with their corresponding wild type B73 (n = 7 – 10). Different letters above the bars indicate statistically significant differences between genotypes as determined by a one-way analysis of variance (ANOVA), followed by estimated marginal means *post hoc* tests (Tukey-adjusted $P \leq 0.05$). FW, fresh weight

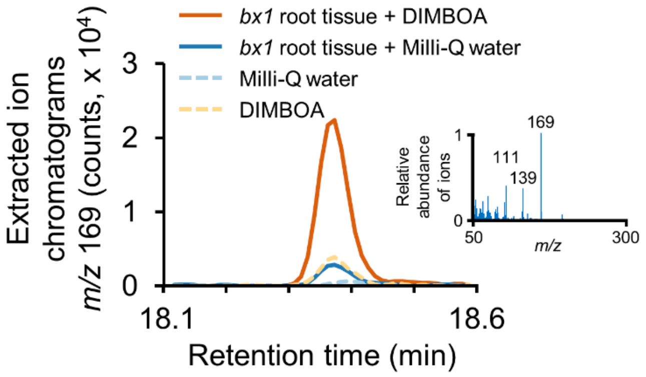


**Figure S6:** 5-Methoxy-2-Nitrophenol (5M2NP) accumulates in ground root tissue of *bx1* mutants supplemented with DIMBOA. Representative GC-MS extracted ion chromatograms of a diagnostic electron ionization ion *(m/z* 169) of 5M2NP, identified in vials loaded with 50 µg DIMBOA either or not in presence of *c*. 100 mg ground crown roots of 16-d-old *bx1* mutants and incubated for 6.1 h (samples with *bx1* root tissue) to 10 h (control samples without *bx1* root tissue), respectively, at room temperature, along with their respective controls. Inset: reference electron ionization mass spectrum of the 5M2NP peak from the *bx1* root tissue + DIMBOA sample. The mass-to-charge ratio of the three most abundant ions is included.

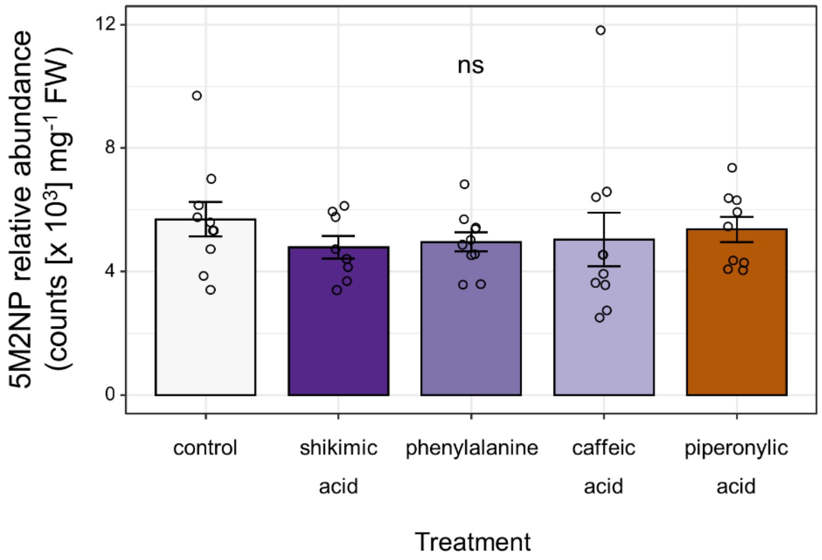


**Figure S7:** The crown root 5-Methoxy-2-Nitrophenol (5M2NP) content of maize (*Zea mays*) seedlings is not significantly altered by various pharmacological treatments aimed at modifying the metabolic flux through the phenylpropanoid pathway. Wild type W22 plants were germinated and grown under soil-free conditions while the roots were repeatedly supplemented with either a precursor (shikimic acid or phenylalanine), an intermediate (caffeic acid) or an inhibitor (piperonylic acid) of the phenylpropanoid pathway. Bars indicate mean (± SEM) amounts of 5M2NP in crown roots of 23-d-old W22 plants subjected to the various pharmacological treatments (n = 8 – 10). The effect of the pharmacological treatments on the amount of 5M2NP in roots was not statistically significant (ns; $P > 0.6$), as determined by a one-way analysis of variance (ANOVA).

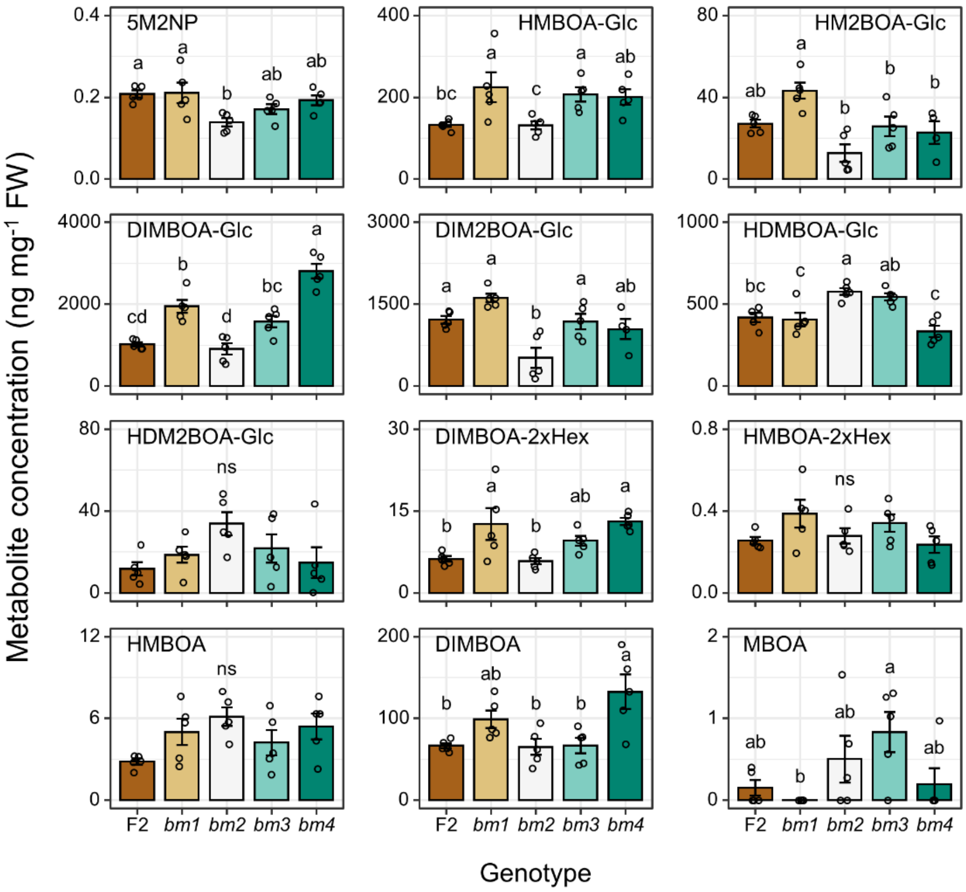


**Figure S8:** Concentrations of 5-Methoxy-2-Nitrophenol (5M2NP) and eleven benzoxazinoids in the crown roots of *brown-midrib* mutants (*bm1-4*) and their corresponding wild type F2. Bars indicate mean (± SEM) amounts of each metabolite in crown roots of 16-d-old seedlings (n = 5). In each panel, different letters above the bars indicate statistically significant differences between genotypes as determined by a one-way analysis of variance (ANOVA), followed by estimated marginal means *post hoc* tests (Tukey-adjusted $P \leq 0.05$). ns, not significant (ANOVA; $P > 0.05$)

(a)

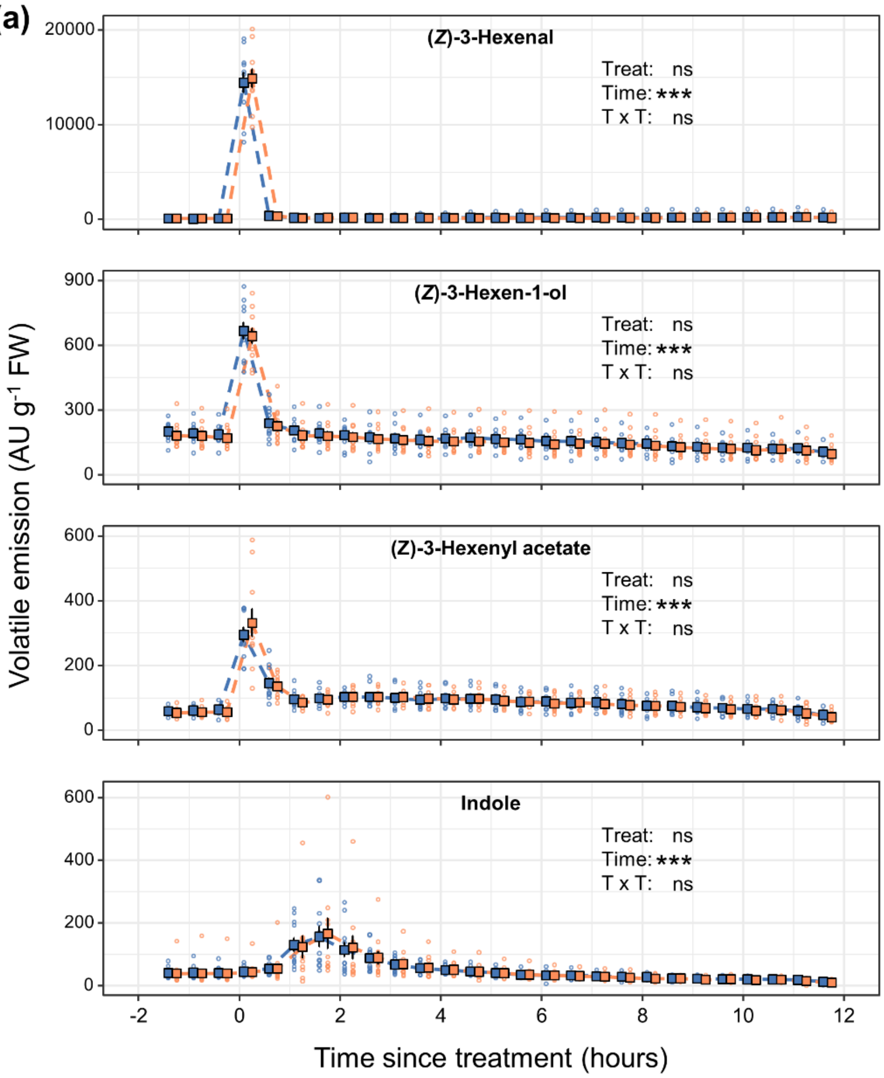

(Z)-3-Hexenal
(Z)-3-Hexen-1-ol
(Z)-3-Hexenyl acetate
Indole
Treat: ns
Time: ***
T x T: ns
Volatile emission (AU g-1 FW)
Time since treatment (hours)

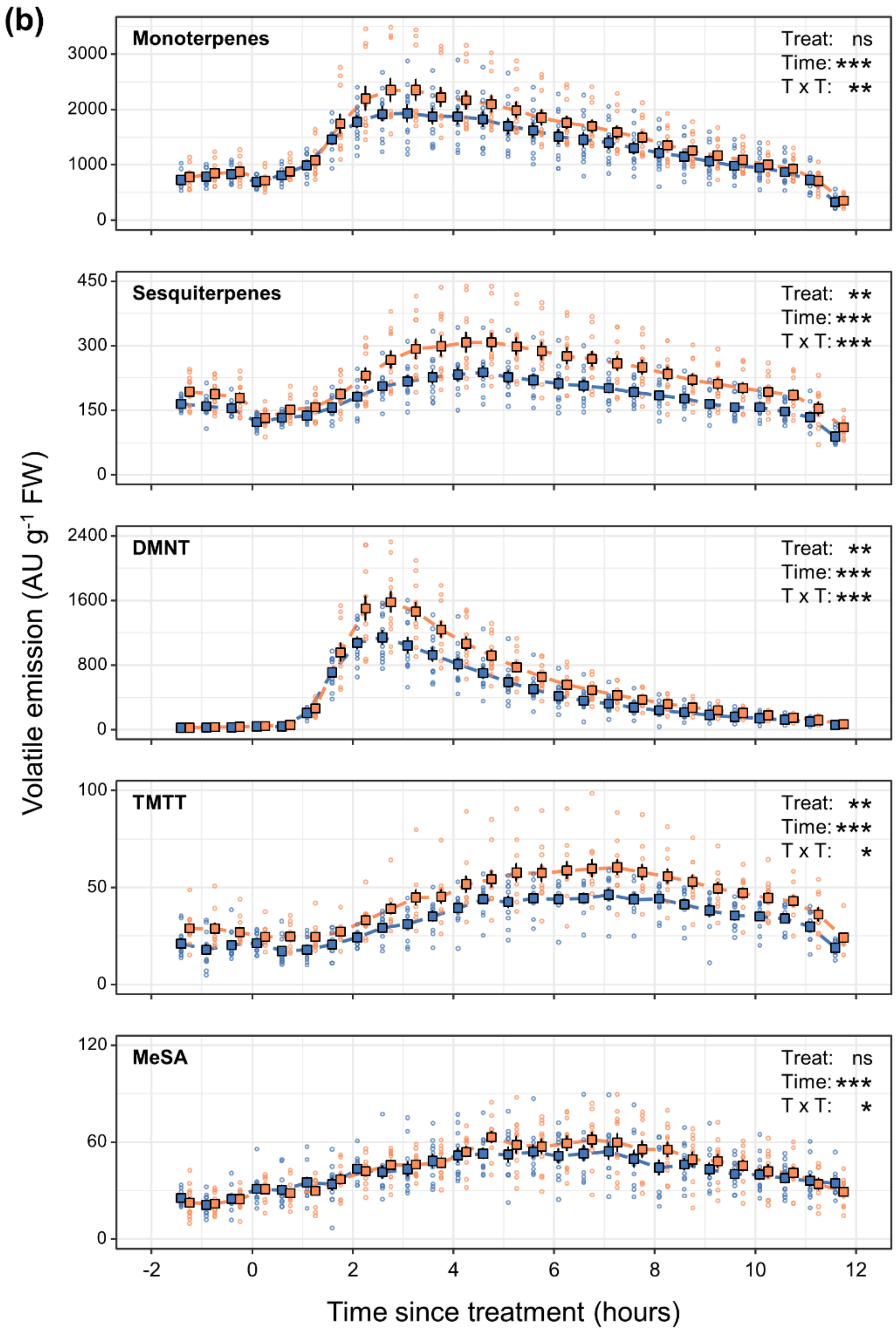

(b)
Monoterpenes
Treat: ns
Time: ***
T x T: **
Sesquiterpenes
Treat: **
Time: ***
T x T: ***
DMNT
Treat: **
Time: ***
T x T: ***
TMTT
Treat: **
Time: ***
T x T: *
MeSA
Treat: ns
Time: ***
T x T: *
Volatile emission (AU g-1 FW)
Time since treatment (hours)

**Figure S9:** 5-Methoxy-2-Nitrophenol (5M2NP) enhances the wound-induced emission of maize volatile organic compounds (VOCs). The figure shows time-resolved emission profiles of the main wound and defense-associated VOCs from maize (*Zea mays*) *bx1* mutants following either a leaf wound + 5M2NP treatment (in orange) or a wound + water treatment (in blue). The third leaf (L3) of 16-d-old *bx1* seedlings was wounded four times, perpendicular to but not crossing the midrib, using a hemostat. Wounds were immediately supplemented with 20 µl of a 5M2NP solution (7.5 ng 5M2NP $\mu l^{-1}$) or Milli-Q water (0.75% v/v dichloromethane) as a control. Squares depict mean (± SEM) normalized emission values of: (a) (*Z*)-3-hexenal ($C_6H_{11}O^+$), (*Z*)-3-hexen-1-ol ($C_6H_{13}O^+$), (*Z*)-3-hexenyl acetate ($C_8H_{15}O_2^+$), indole ($C_8H_8N^+$), and (b) monoterpenes ($C_{10}H_{17}^+$), sesquiterpenes ($C_{15}H_{25}^+$), DMNT ($C_{11}H_{19}^+$), TMTT ($C_{16}H_{27}^+$) and methyl salicylate (MeSA; $C_8H_9O_3^+$) for each treatment, at indicated time points (n = 11). Regarding the post-treatment data, black asterisks denote statistically significant effects of plant treatment (Treat), time since treatment (Time) and/or their interaction (T x T) on the emission of each VOC, as determined by a multifactorial repeated measures analysis of variance on aligned rank-transformed data (RM-ANOVA-ART): *** for $P < 0.001$; ** for $P < 0.01$; * for $P < 0.05$; ns, not significant for $P > 0.1$. Data from the final sampling time point were the first to be collected in the artificial night and are shown here solely for reference purposes (maize plants emit hardly any terpenoids during the night), i.e., they were not included in any statistical analysis. Detailed results from the statistical analysis, including effects of plant treatment at individual time points, are listed in Table S2.

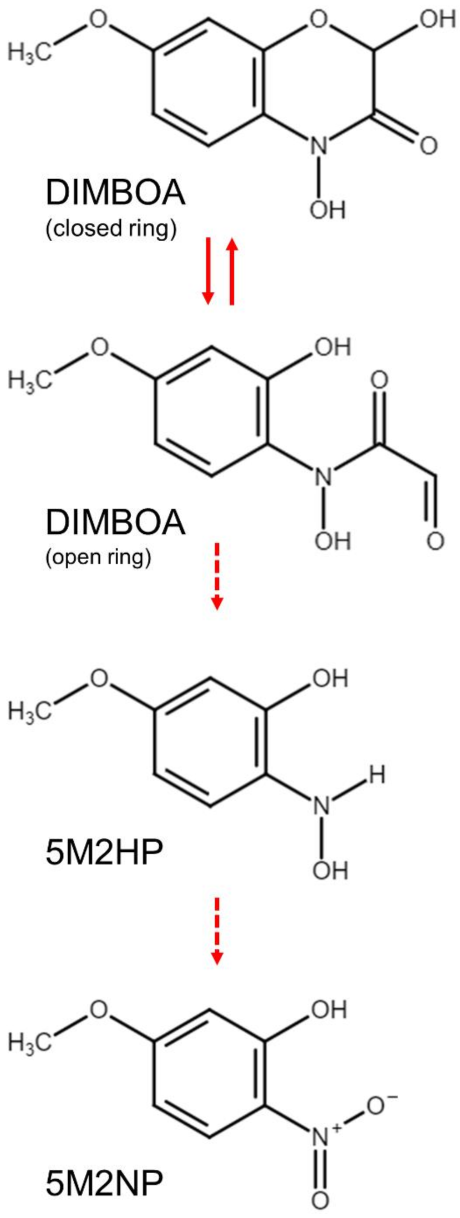


**Figure S10:** Hypothetical metabolic route from DIMBOA to 5-Methoxy-2-Nitrophenol (5M2NP). Breakage of the nitrogen-carbonyl bond in open-form-DIMBOA and subsequent oxidation of the putative 5-Methoxy-2-Hydroxyaminophenol (5M2HP) intermediate could yield 5M2NP. Solid arrows represent known chemical reactions, whereas dashed arrows represent one or more uncharacterized, putative chemical transformations.

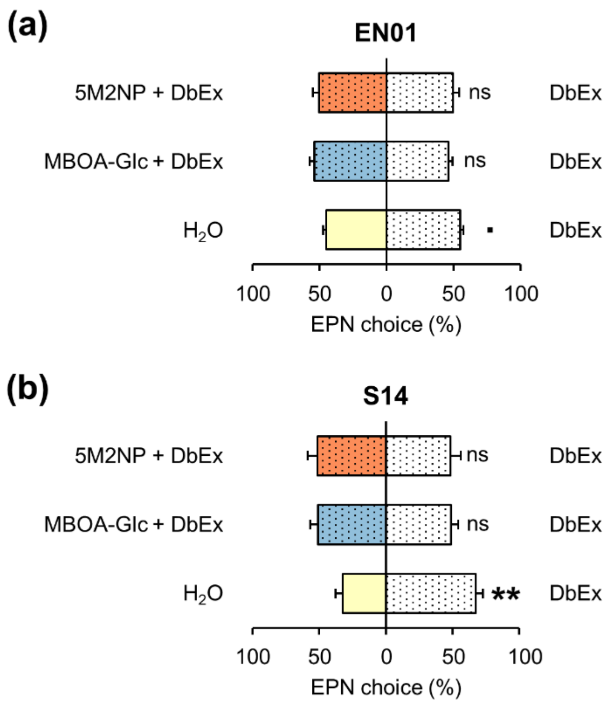


**Figure S11:** At a physiologically relevant dose, 5-Methoxy-2-Nitrophenol (5M2NP) does not significantly alter the foraging behavior of *Heterorhabditis bacteriophora* entomopathogenic nematodes (EPNs) in Petri dish-based choice assays. Bars indicate mean (± SEM) proportions of EPNs residing in sections with (1) water (as a control) or exudates from *bx1/W22*-fed *D. balteata* larvae (referred to as 'DbEx'), (2) DbEx or DbEx supplemented with 330 ng MBOA-Glc, and; (3) DbEx or DbEx supplemented with 25 ng 5M2NP (n = 20 choice situations with 100 EPNs each). All treatment solutions contained 0.5% (v/v) ethanol. Black asterisks denote statistically significant differences between treatments according to a one-sample *t*-test or a one-sample Wilcoxon signed rank test based on the percentage of EPNs that chose for either of the treatments (*** for $P < 0.001$; ** for $P < 0.01$; * for $P < 0.05$; . for $P < 0.1$.; ns, not significant for $P > 0.1$). EPN choice assays were performed with (a) the benzoxazinoid-susceptible strain EN01 and (b) the benzoxazinoid-resistant strain S14. See Methods S9 for experimental details about EPN preference assays.

**Table S1:** qRT-PCR primer specifications

| Gene | Full name | Gene Identifier | Forward Primer (5’ → 3’) | Reverse Primer (5’ → 3’) | Primer reference |
|---|---|---|---|---|---|
| *AOS1* | *Allene oxide synthase 1* | GRMZM2G067225 | ACAAGGTGGAGAAGAAGGAC | GTCGTTGAGCTTGTTGAACT | Huffaker et al. (2013) |
| *AOC1* | *Allene oxide cyclase 1* | GRMZM2G077316 | CTTCTACTTCGGAGACTACGG | TCAGTTGGTGTAGTTGTCGAG | This study |
| *SerPIN* | *Serine proteinase inhibitor* | GRMZM5G815098 | GACCTCAGGCGTCGTTACTC | TGCAGCAAATATAGCCAACG | Huffaker et al. 2013 |
| *Cyst/CC9* | *Cystatin proteinase inhibitor* | Zm00001d049111 | CAAGGAGCACAACAGGCAGA | GGACATGAGCTGGCGATTTT | Ton et al. (2007) |
| *LOX10* | *Lipoxygenase 10* | Zm00001d053675 | ATCCTCAGCATGCATTAGTCC | AGTCTCAAACGTGCCTCTTGT | Vaughan et al. (2014) |
| *TPS10* | *Terpene synthase 10* | Zm00001d024486 | TGACAGCCTTGATCACCGTA | AGTTCATCCCCACACGAATC | This study |
| *CYP92C5* | *Cytochrome P450 monooxygenase 92C5* | Zm00001d018839 | AGGGGTTCAAGCGGAAGATG | AGGTCAGTCGCCACAAACTC | This study |
| *IGL* | *Indole-3-glycerol phosphate lyase* | Zm00001d034453 | GAGCTGGTGCTGCTCACA | GCGCGTGGACCTGTAACT | This study |
| *Bx1* | *Benzoxazinless 1* | Zm00001d048709 | GCTTCGTCTACCTGGTGAGC | ACAGCAACGGGCTTGTTAGT | This study |
| *Bx10/11* | *Benzoxazinless 10/ Benzoxazinless 11* | GRMZM2G311036/ GRMZM2G336824 | CAGCAGGTGGTGGTGATAAT | AGCGCCAGACTCACAAAGG | Meihls et al. (2013) |
| *Bx14* | *Benzoxazinless 14* | GRMZM2G127418 | GAAAGCCGCTTCTTGATGCC | GGAACATATTGCCCGCAACG | Maag et al. (2016) |
| *Ubi1* | *Ubiquitin* | Zm00001d015327 | TAAGCTGCCGATGTGCCTGCG | CTGAAAGACAGAACATAATGAGCACAG | Bommert et al. (2005) |

**Table S2:** Results from statistical analysis of maize headspace volatile organic compounds data, presented in Figs. 4 and S9.

| | (*Z*)-3-hexenal | (*Z*)-3-hexen-1-ol | (*Z*)-3-hexenyl acetate | GLVs | Indole | Monoterpenes | Sesquiterpenes | DMNT | TMTT | Homoterpenes | MeSA |
|---|---|---|---|---|---|---|---|---|---|---|---|
| **Pre-treatments (-80 ≤ time [min] ≥ -20)**<br>*P*-values from multifactorial repeated measures analysis of variance on aligned rank transformed data (RM-ANOVA-ART) | | | | | | | | | | | |
| **Treatment** | 0.40644 | 0.21910 | 0.15824 | 0.42640 | 0.10711 | 0.55487 | 0.11428 | 0.05639 | 0.01420 | 0.01743 | 0.64314 |
| **Time** | 0.19917 | 0.01027 | 0.03672 | 0.68442 | 0.91244 | 1.18e-11 | 4.18e-05 | 6.23e-08 | 0.07508 | 2.88e-05 | 0.14757 |
| **T x T** | 0.21586 | 0.68241 | 0.70289 | 0.38451 | 0.82871 | 0.68161 | 0.61222 | 0.26141 | 0.05567 | 0.01361 | 0.34317 |
| **Post-treatments (10 ≤ time [min] ≥ 670; i.e. until the end of the light period)**<br>*P*-values from multifactorial repeated measures analysis of variance on aligned rank transformed data (RM-ANOVA-ART) | | | | | | | | | | | |
| **Treatment** | 0.34663 | 0.33370 | 0.89509 | 0.94335 | 0.67746 | 0.26711 | 0.00914 | 0.00356 | 0.00129 | 0.00296 | 0.38322 |
| **Time** | < 2e-16 | < 2e-16 | < 2e-16 | < 2e-16 | < 2e-16 | < 2.22e-16 | < 2.22e-16 | < 2e-16 | < 2.22e-16 | < 2.22e-16 | < 2e-16 |
| **T x T** | 0.99975 | 0.98041 | 0.91667 | 0.99999 | 0.99885 | 0.00562 | 2.25e-10 | < 2e-16 | 0.04664 | < 2.22e-16 | 0.01449 |
| **Individual time points**<br>non-adjusted *P*-values, either from independent two-sample Student's *t*-test, Welch's *t*-test or a Wilcoxon rank-sum test | | | | | | | | | | | |
| **Time (min)** | | | | | | | | | | | |
| -80 | 0.27032 | 0.19319 | 0.30575 | 0.43846 | 0.13299 | 0.46426 | 0.03606 | 0.51903 | 0.03360 | 0.12384 | 0.41989 |
| -50 | 0.39300 | 0.59468 | 0.17131 | 0.95880 | 0.17131 | 0.42011 | 0.04867 | 0.04836 | 0.02331 | 0.01923 | 0.77181 |
| -20 | 0.89766 | 0.15127 | 0.22205 | 0.21695 | 0.10142 | 0.57234 | 0.08272 | 0.08954 | 0.02331 | 0.01038 | 0.97365 |
| 10 | 0.76488 | 0.64405 | 0.45251 | 0.76712 | 0.13299 | 0.68347 | 0.29717 | 0.26815 | 0.17458 | 0.18355 | 0.98040 |
| 40 | 0.68596 | 0.67254 | 0.59976 | 0.63198 | 0.08795 | 0.40092 | 0.05748 | 0.00457 | 0.01577 | 0.00354 | 0.89766 |
| 70 | 0.69935 | 0.17131 | 0.25346 | 0.38478 | 0.47785 | 0.40779 | 0.05869 | 0.08199 | 0.02807 | 0.05237 | 0.21863 |
| 100 | 0.65218 | 0.36531 | 0.65111 | 0.87767 | 0.69935 | 0.40092 | 0.04578 | 0.06280 | 0.01766 | 0.05753 | 0.55964 |
| 130 | 0.84700 | 0.73960 | 0.96303 | 0.89788 | 0.74766 | 0.29999 | 0.02959 | 0.02598 | 0.00478 | 0.02424 | 0.67408 |
| 160 | 0.74766 | 0.43846 | 0.69935 | 0.84041 | 0.43846 | 0.24265 | 0.01430 | 0.01393 | 0.00663 | 0.01263 | 0.21695 |
| 190 | 0.74766 | 0.65218 | 0.81182 | 0.86297 | 0.79694 | 0.19319 | 0.00998 | 0.01246 | 0.00663 | 0.01100 | 0.17131 |
| 220 | 0.79694 | 0.83321 | 0.70060 | 1.00000 | 0.56189 | 0.27032 | 0.01867 | 0.02801 | 0.03939 | 0.02539 | 0.81498 |
| 250 | 0.65218 | 0.33165 | 0.75001 | 0.74766 | 0.51903 | 0.33165 | 0.01589 | 0.04230 | 0.01923 | 0.03581 | 0.71237 |
| 280 | 0.94873 | 0.29999 | 0.99861 | 0.56189 | 0.51903 | 0.29999 | 0.01581 | 0.04921 | 0.05868 | 0.04183 | 0.06408 |
| 310 | 0.74766 | 0.24265 | 0.65231 | 0.74766 | 0.74766 | 0.08795 | 0.00912 | 0.05826 | 0.01291 | 0.04477 | 0.41770 |
| 340 | 0.74766 | 0.24265 | 0.97827 | 0.74766 | 0.51903 | 0.25702 | 0.01128 | 0.05381 | 0.01982 | 0.03997 | 0.60820 |
| 370 | 0.84700 | 0.21695 | 0.56474 | 0.60632 | 0.93546 | 0.24265 | 0.01692 | 0.04809 | 0.00140 | 0.03384 | 0.25261 |
| 400 | 0.84700 | 0.27032 | 0.92017 | 0.60632 | 0.60632 | 0.15127 | 0.01250 | 0.04300 | 0.00186 | 0.02747 | 0.23424 |
| 430 | 0.60632 | 0.29999 | 0.61619 | 0.79694 | 0.51903 | 0.26240 | 0.01095 | 0.05337 | 0.01073 | 0.03478 | 0.48067 |
| 460 | 1.00000 | 0.19319 | 0.73422 | 0.65218 | 0.51886 | 0.20607 | 0.00954 | 0.04285 | 0.00775 | 0.02549 | 0.42129 |
| 490 | 0.89766 | 0.36531 | 0.95032 | 0.79694 | 0.17397 | 0.35524 | 0.02067 | 0.07028 | 0.01838 | 0.04809 | 0.11256 |
| 520 | 0.84700 | 0.51903 | 0.72887 | 1.00000 | 0.98229 | 0.41606 | 0.02004 | 0.08497 | 0.01376 | 0.05272 | 0.62503 |
| 550 | 0.89766 | 0.33165 | 0.74596 | 0.74766 | 0.17131 | 0.41306 | 0.01111 | 0.06543 | 0.01284 | 0.03613 | 0.38257 |
| 580 | 1.00000 | 0.36531 | 0.60787 | 0.79694 | 0.82317 | 0.32374 | 0.01061 | 0.06221 | 0.00286 | 0.03162 | 0.28258 |
| 610 | 0.74766 | 0.29999 | 0.56189 | 0.69935 | 0.22549 | 0.60569 | 0.01953 | 0.12368 | 0.01783 | 0.06987 | 0.73275 |
| 640 | 0.94873 | 0.29999 | 0.59024 | 0.84700 | 0.59685 | 0.55243 | 0.01140 | 0.18452 | 0.01039 | 0.09803 | 0.49427 |
| 670 | 0.97254 | 0.35160 | 0.30776 | 0.56189 | 0.13636 | 0.95536 | 0.20468 | 0.37328 | 0.13633 | 0.28730 | 0.66210 |
| 700 | 0.65218 | 0.49084 | 0.24265 | 0.51903 | 0.29454 | 0.55888 | 0.01194 | 0.41209 | 0.00663 | 0.23233 | 0.14978 |

**Methods S1**

**5M2NP quantification by means of UHPLC-MS/MS**

Concentrations of 5M2NP in sampled plant tissues were determined by UHPLC-MS/MS, using an Acquity I-Class UPLC system (Waters Corp., Milford, MA, USA) coupled to a Xevo TQ-XS triple quadrupole mass spectrometer (Waters) through an atmospheric pressure chemical ionization (APCI) source. The entire system was controlled by MassLynx software v4.2 (Waters). Plant treatment and sample preparation details were as follows. The second leaf of 14-d-old B73 seedlings was wounded six times, perpendicular to but not crossing the midrib, using a hemostat. Ten minutes later, the wounded leaf was cut off from the plant, flash frozen in liquid nitrogen and stored at -80°C until further processing (n = 4). Corresponding leaves from untreated plants were sampled and used as controls. Three 100 mg aliquots of ground frozen homogenized leaf material were produced from each biological replicate. The first aliquot was incubated at room temperature for 20 minutes, thus allowing the tissue to thaw. The second aliquot was incubated at 40°C for 20 minutes. As a control, the third aliquot was not subjected to a secondary treatment (i.e., kept at -80°C). Subsequently, 500 µl of 100% methanol was added to each sample, and vortexed for 10 sec. Samples were centrifuged at 17,000 g for 2 min, and 200 µl of the supernatant was transferred to glass vials for analysis. For this, 3 µL was injected into a Cortecs C18+ column (50 x 2.1mm, 1.6 µm particle size, Waters). The mobile phases consisted of Milli-Q water (A) and methanol (B). The flow rate was 0.4 mL $min^{-1}$. A gradient program starting at 10% phase B and linearly increasing to 52% phase B in 6 min was applied, then the proportion of B was rapidly increased to 100% in 0.5 min and maintained for 2.0 min before re-equilibrating the column at 10% B for 3 min. The temperature of the column was kept at 25°C. Under these conditions, 5M2NP eluted at 5.83 min. The APCI source was operated in negative ionization mode using the following parameters: corona current 15 µA, probe temperature 425°C, source temperature 150°C, desolvation gas flow 900 L $h^{-1}$, nebulizer gas flow 4.0 bars, cone gas flow 350 L $h^{-1}$. The mass spectrometer was operated in the multiple reaction monitoring (MRM) mode using the following quantitative (Q) and qualitative (q) transitions: 168/153 (Q), 168/125 (q1), 168/123 (q2), and 168/95 (q3). The cone voltage was set to 20 V for all MRM transitions and the collision energies to 14, 20, 20, and 25 V for transitions Q, q1, q2, and q3, respectively. Quantification was achieved by external calibration using calibration points at 2.5, 5, 20, 50 and 200 ng $mL^{-1}$. A linear regression weighted by 1/x was applied. TargetLynx XS software (Waters) was used for data analysis.

**Methods S2**

**Gene expression analysis**

Transcript abundances of selected marker genes in harvested leaf sections were determined by qRT-PCR following the procedures described by Schimmel et al. (2017) and Escobar-Bravo et al. (2022), with minor modifications. Briefly, total RNA was isolated using the GeneJET Plant RNA Purification Kit (Thermo Fisher Scientific, Waltham, MA, USA), with the method scaled up to process up to 150 mg of leaf tissue per sample. DNAse treatment and cDNA synthesis were carried out using the PrimeScript RT reagent Kit with gDNA eraser (Perfect Real Time; TaKaRa Bio Inc., Kusatsu, Japan). For gene expression analysis, 2 µl of diluted cDNA, i.e., the equivalent of 10 ng total RNA, served as template in a 10 µl qRT-PCR using ORA SEE qPCR Green ROX L Mix (highQu GmbH, Kraichtal, Germany) on a 384-well Applied Biosystems QuantStudio 5 Real-Time PCR system (Thermo Fisher Scientific), according to the manufacturer's instructions. Normalized expression (NE) values were calculated with the ΔCt method: $NE = (1/(PE_{target}\string^Ct_{target}))/(1/(PE_{reference}\string^Ct_{reference}))$ in which PE stands for 'primer efficiency' and Ct for 'cycle threshold' (Alba et al., 2015). PEs were calculated by fitting a linear regression on the Ct values of a standard cDNA dilution series. *Ubiquitin* was used as the reference gene. Target genes, their identifiers, primer sequences and references are listed in Table S1.

**Methods S3**

**Phytohormone analysis**

Phytohormone concentrations in sampled leaf sections were determined by UHPLC-MS/MS, using an Acquity UPLC I-Class system (Waters) connected to a QTRAP 6500+ mass spectrometer (SCIEX, Framingham, MA, USA) as in Glauser et al. (2014) and Schimmel et al. (2017), with minor modifications. In short, 1 ml of ethyl acetate:formic acid (99.5:0.5 v/v), spiked with $d_6$-ABA, $d_5$-IAA, $d_5$-JA, $^{13}C_6$-JA-Ile and $d_6$-SA at 100 ng $ml^{-1}$, was added to frozen leaf powder (100-150 mg) inside a 2 ml microcentrifuge tube, which also contained three 2.8 mm zirconium oxide beads, and vortexed for 10 sec. Hormones were then extracted in a mixer mill (Retsch MM300; Retsch GmbH, Haan, Germany) at 30 Hz for 3 min. Samples were centrifuged at 14,000 g for 4 min and the supernatant was transferred to new tubes. The extraction step was repeated with the pellet and 0.5 ml of ethyl acetate:formic acid (99.5:0.5% v/v) without internal standards. Both supernatants were combined and evaporated to dryness on a centrifugal concentrator (CentriVap Centrifugal Concentrator, Labconco, Kansas City, MO, USA) at 35°C. The residue was re-suspended in 200 µl of 50% methanol and after two consecutive centrifugation steps (at 14,000 g for 3 min) to exclude plant particles, 120 µl of the supernatant was transferred to glass vials for analysis. For this, 1.5 µl of extract was injected onto an Acquity UPLC BEH C18 column (50 x 2.1 mm, 1.7 µm particle size; Waters). Mobile phase A consisted of $H_2O$:formic acid (99.95:0.05% v/v) and mobile phase B consisted of acetonitrile:formic acid (99.95:0.05% v/v). A gradient from 5-65% B in 6.5 min was applied, followed by column washing at 100% B and equilibration at 5% B for 2 min. The flow rate was set to 0.4 ml $min^{-1}$ and the column temperature to 35°C. The mass spectrometer was operated in electrospray negative ionization using the multiple reaction monitoring (MRM) mode. A six-point calibration curve (0.02, 0.1, 0.5, 5, 20, and 100 ng $ml^{-1}$, containing all isotopically labelled standards at 5 ng $ml^{-1}$) was used for quantification. Linear regressions weighted by 1/x were applied. Analyst v.1.7.1 was used to control the instrument and for data processing.

**Methods S4**

**Caterpillar preference assays**

Choice assays with *S. littoralis* and *S. frugiperda* larvae were adapted from Glauser et al. (2011) and Rharrabe et al. (2014). We monitored the distribution over time of groups of five naïve caterpillars given a choice between two differently treated cubes of a soy-wheat germ-based artificial diet (AD) placed in a Petri dish (9 cm diameter; Greiner Bio-One, Kremsmünster, Austria) at a 5 cm distance from each other. AD cubes (c. 5 x 10 x 5 mm) were supplemented with either a 5M2NP solution or Milli-Q water (0.9% v/v ethanol) as a control, applied at 1.0 µl $mg^{-1}$ AD. Two assays were performed in parallel: one with 1.0 ng 5M2NP $mg^{-1}$ AD and another with 2.5 ng 5M2NP $mg^{-1}$ AD. We used third-instar larvae and starved them for 24 h prior to the start, i.e. their release into the center of a Petri dish. The number of larvae residing in the quarters containing the AD cubes were counted at 10 min, 20 min, 30 min, 60 min, 120 min and 960 min after the start of the experiment. The assays were carried out in a laboratory room (without windows) at RT under artificial light until the 120 min time point, after which caterpillars were left overnight in the dark until the 960 min time point. The amounts of 5M2NP for AD complementation were determined based on the 5M2NP concentration found in maize leaves and represent an average and a high-end value, respectively (Fig. S2). The AD itself does not emit 5M2NP, as determined by SPME-GC-MS. These experiments were performed with 20 Petri dishes (choice situations), containing five larvae each, per caterpillar species and treatment combination (n = 20).

**Methods S5**

**Rootworm preference assays**

Choice assays with *D. balteata* and *D. v. virgifera* larvae were adapted from larval-escape experiments described by Huang et al. (2017). We assessed the extent to which supplementation of 5M2NP to the rhizosphere of *bx1* plants affected the host rejection rate of *Diabrotica* larvae. To do so, 14-d-old *bx1* seedlings were transplanted individually into L-shaped glass pots (height 11 cm, diameter 5 cm) with a horizontal connector (length 6.5 cm, diameter 2.9-3.2 cm) at a height of 0.5 cm (Verre et Quartz Technique SA, Neuchâtel, Switzerland). Two days later, two holes (depth 10 cm, diameter 0.5 cm) were pierced into the rhizosphere on opposite sides of the stem and used to introduce 5 ml of either a 5M2NP solution or autoclaved tap water (0.0025% v/v ethanol) as a control. After a 10 min incubation period, five *bx1/W22*-reared third-instar larvae were released at the entrance of the horizontal access port of each pot. The larvae were starved for 1 h prior to the start of the assays. Once released, larvae were free to move into the rhizosphere and start feeding, or alternatively, to turn around and move away from the host plant, in which case they were trapped in plastic cups. The number of host-rejecting larvae was counted after 20 min. We used 5M2NP solutions with two distinct concentrations such that the total amount of 5M2NP added to the rhizosphere of each plant was either 25 ng or 125 ng. These amounts of 5M2NP for complementation were determined based on the average 5M2NP concentration found in crown roots of WT maize (0.1 ng $mg^{-1}$ FW; Fig S2), a conservative estimate of *D. v. virgifera*'s root consumption (250 mg FW $day^{-1}$ by five larvae; Zhang et al., 2019a), and the average mass of the total root system of 16-d-old *bx1* seedlings (*c.* 1100 mg). Using these values, we reasoned that 5M2NP produced in roots of WT plants attacked by five rootworms readily amounts to 25 ng $day^{-1}$. Hence, the highest amount of 5M2NP added reflects a more severe rootworm attack. These experiments were performed in three or four blocks (experimental replicates) in time, in total with 37 to 58 plants (choice situations), with five larvae each, per rootworm species and treatment (n = 37 – 58). Note that for both species, all three treatments were included in three of the experimental replicates, while a fourth replicate was performed with only two treatments, i.e. the control treatment and one of the 5M2NP treatments. Consequently, possible effects of the two 5M2NP treatments on rootworm behavior were analyzed separately.

**Methods S6**

**Caterpillar performance assays**

No-choice performance assays with *S. littoralis* and *S. frugiperda* larvae were modified from Glauser et al. (2011). That is, using 1.25 oz. plastic cups (Frontier Agricultural Sciences), we monitored the weight over time of individual naïve caterpillars feeding on AD supplemented with either a 5M2NP solution or Milli-Q water (0.9% v/v ethanol) as a control, applied at 1 μl $mg^{-1}$ AD. Two concentrations of 5M2NP were tested in parallel: 1.0 ng 5M2NP $mg^{-1}$ AD and 2.5 ng 5M2NP $mg^{-1}$ AD. We used pre-weighed second-instar larvae and recorded their weight and replaced their diet every other day until experiments were terminated at day 12, i.e. when further *ad libitum* feeding for *S. littoralis* larvae could no longer be guaranteed. Treatment solutions were freshly prepared on each diet-replacement day. Except during weighing and diet replacement, larvae were kept in the dark at RT. These assays were performed with 30 larvae per caterpillar species and treatment (n = 30).

**Methods S7**

**Rootworm performance assays**

No-choice performance assays with *D. balteata* and *D. v. virgifera* were based on methods presented in Erb et al. (2011). Here, we determined the weight gain of naïve *Diabrotica* larvae feeding from *bx1* plants supplemented with either a 5M2NP solution or autoclaved tap water (0.0025% v/v ethanol) as a control. For this, 5 ml of a treatment solution was first pipetted onto the soil (rhizosphere) of 16-d-old *bx1* seedlings, after which each plant was infested with five pre-weighed, *bx1/W22*-reared, third-instar larvae. We added 5M2NP solutions with three distinct concentrations such that the daily total amount of 5M2NP added to the rhizosphere of each plant was either 2.5 ng, 25 ng or 125 ng. Plants were supplemented with 5 ml of treatment solution for four consecutive days, after which the larvae were recovered and weighed. Note that independent of the treatment and rootworm species, we were unable to consistently recover all larvae. These assays were performed with 12 plants per rootworm species and treatment (n = 12).

**Methods S8**

**Plant infestations with herbivores**

Maize infestations with caterpillars and rootworms were performed as described by Glauser et al. (2011) and Huang et al. (2017). In a first experiment, 21-d-old Delprim plants were infested with either twelve second-instar or three third-instar *S. littoralis* larvae. Uninfested plants served as controls. Two days later, leaves, crown roots and primary roots were collected (separately), flash frozen in liquid nitrogen and stored at -80°C until analysis by SPME-GC-MS. This experiment was performed with ten plants per treatment (n = 10). In a second experiment, 14-d-old B73 plants were infested with twelve second-instar *S. frugiperda* larvae. Uninfested plants served as controls. Two days later, leaves and crown roots were collected, flash frozen in liquid nitrogen and stored at -80°C until analysis by SPME-GC-MS. This experiment was performed with seven plants per treatment (n = 7). In a third experiment, 12-d-old Delprim plants were infested with either six W22-reared, third-instar *D. balteata* or *D. v. virgifera* larvae. Uninfested plants served as controls. Four days later, leaves, crown roots and primary roots were collected, flash frozen in liquid nitrogen and stored at -80°C until analysis by SPME-GC-MS. This experiment was performed in two blocks (experimental replicates) in time, in total with twelve plants per treatment (n = 12).

**Methods S9**

**EPN preference assays**

Petri dish-based EPN choice assays were conducted as in Robert et al. (2017), with some modifications. We generated Petri dishes (9 cm diameter; Greiner Bio-One) with a 5 mm layer of 0.5% agarose (Sigma Aldrich) containing three circular wells along the plate diameter: a large one in the center (10 mm diameter) and two smaller ones (5 mm diameter) on either side at a distance of 2.5 cm from the center. The small wells were each filled with 50 µl of a test solution, followed by the release of 100 EPNs suspended in 100 µl autoclaved tap water into the central well. Petri dishes were then kept in the dark at RT and the number of EPNs residing in the quarters containing the wells with test solutions were counted 24 h later. Except for the negative control (50 µl autoclaved tap water), all test solutions contained 45 µl of exudates from *bx1/W22*-fed *D. balteata* larvae (referred to as "DbEx") to trigger EPN foraging behavior. DbEx was collected from third-instar larvae using autoclaved tap water (50 µl per larva) and stored at -20°C until use. Test solutions further contained 5 µl of either a 5M2NP solution, an MBOA-Glc solution, or autoclaved Milli-Q water (0.5% v/v ethanol) as a control. The total amounts of 5M2NP and MBOA-Glc used for DbEx complementation were 25 ng and 330 ng, respectively, per well. The final MBOA-Glc concentration corresponds to that found on the skin of third-instar *D. v. virgifera* larvae, which is sufficient to repel benzoxazinoid-susceptible EPNs (Robert et al., 2017). We surveyed the foraging behavior of *H. bacteriophora* nematodes from the benzoxazinoid-susceptible strain ENO1 (Zhang et al., 2019b; referred to as the Andermatt strain in Robert et al., 2017) and the benzoxazinoid-resistant strain S14 (Zhang et al., 2019b) in the following two-choice assays: (i) water vs. DbEx, (ii) DbEx vs. DbEx + 5M2NP, and (iii) DbEx vs. DbEx + MBOA-Glc. These assays were performed with 20 Petri dishes (choice situations), containing 100 EPNs each, per nematode strain and treatment combination (n = 20). Nematode positions were recorded without knowledge of treatment allocations (blinded observations).
